# Divergent Directed Evolution of a TetR-type Repressor Towards Aromatic Molecules

**DOI:** 10.1101/2022.06.12.495817

**Authors:** Mohamed A. Nasr, Vincent J.J. Martin, David H. Kwan

## Abstract

Reprogramming cellular behaviour is one of the hallmarks of synthetic biology. To this end, prokaryotic allosteric transcription factors (aTF) have been repurposed as versatile tools for processing small molecule signals into cellular responses. Expanding the toolbox of aTFs that recognize new inducer molecules is of considerable interest in many applications. Here, we first establish a resorcinol responsive aTF-based biosensor in *Escherichia coli* using the TetR-family repressor RolR from *Corynebacterium glutamicum*. We then perform an iterative walk along the fitness landscape of RolR to identify new inducer specificities, namely catechol, methyl catechol, caffeic acid, protocatechuate, L-DOPA, and the tumour biomarker homovanillic acid. Finally, we demonstrate the versatility of these engineered aTFs by transplanting them into the model eukaryote *Saccharomyces cerevisiae*. This work provides a framework for efficient aTF engineering to expand ligand specificity towards novel molecules on laboratory timescales, which, more broadly, is invaluable across a wide range of applications such as protein and metabolic engineering, as well as point-of-care diagnostics.

## Introduction

Bacteria have evolved complex networks for transcriptional regulation of gene expression to respond to environmental and chemical cues^1–4^. Allosteric transcription factors (aTFs) are regulatory proteins that conditionally bind DNA promoters to regulate gene expression in response to small molecules^5,6^. Bacterial aTFs are frequently employed as tools in synthetic biology for processing biochemical signals to cellular responses. This is in part due to the ease of their heterologous expression as relatively small soluble cytoplasmic proteins, in addition to the wealth of knowledge of aTF structures, DNA recognition sequences (operators), and small molecule inducers^7–9^. As a result, aTF-based biosensors are used in a multitude of applications including but not limited to: screening for environmental contaminants^10,11^, screening metagenomic libraries to identify genes encoding functional enzymes^12^, point-of-care disease diagnostics^13^, as well as controlling synthetic bacterial consortia^14^ and bacterial behaviour *in situ*^15^. As well, aTFs are used in metabolic engineering, which aims to precisely engineer native and heterologous metabolism to produce value-added chemicals^16,17^.

Another attractive aspect of aTF-based biosensors is their tunability. Well-established methods exist for tuning the response curves of genetic circuits such as optimizing promoter and ribosomal binding site (RBS) strength as well as implementing amplifier circuits^18–21^. Tuning the ligand specificity of aTFs however remains a challenging undertaking. aTFs have often evolved to respond to a limited set of inducers, which restricts their use in many applications^22^. The existing repertoire of aTFs could be expanded by continuous bioprospecting^23,24^, creating synthetic chimeras^25,26^, or by directed evolution to alter their ligand specificity^8^. Directed evolution is widely used to introduce beneficial properties to proteins, where rounds of genetic sequence diversification (either random or structure-guided) are followed by variant library screening or selection^27,28^. One of the challenges of aTF directed evolution is maintaining allosteric interdomain signalling for proper protein function^29,30^, with the possible localized unfolding and intrinsic disorder of the DNA-binding domain further complicating this task^31,32^.

In the present work, we report a broadly applicable automated framework for engineering aTFs through the semi-rational directed evolution of the TetR-family member RolR from *Corynebacterium glutamicum*^33,34^. Using combinatorial active site saturation test (CAST)^35^ and iterative saturation mutagenesis (ISM)^36^ methods, which allow for a Cartesian 3-dimensional view of the ligand-binding pocket, we targeted separately and sequentially different regions to generate variants of RolR responsive to aromatic small molecules (Fig. 1a-c). We first target 19 residues in the ligand-binding pocket for mutagenesis, coupled with under-sampling for further variant library size reduction^37^. We then employ the TetA antiporter^38,39^ for dual selection over several variant generations to identify activity towards other aromatic molecules, coupled with an automated screening strategy (Fig 1d-e). RolR-regulated promoters are induced by resorcinol, a molecule with various industrial applications^40^. We start our engineering efforts by targeting catechol biosensing, which is a lignin degradation by-product^41^, and a precursor to the bioproduction of value-added chemicals including muconic acid^42^, as well as the chemical synthesis of fragrances, dyes, and pesticides^43^. We then use the identified catechol-responsive variants as a template for further evolution towards substituted catechol moieties such as lignin degradation by-products (methyl catechol)^41^, pathway intermediates to value-added chemicals (protocatechuate^42^, 2,3-dihydroxybenzoate^44^, L-DOPA^45^, caffeic acid^46^), catecholamine neurotransmitters (epinephrine, dopamine), and 3-*O*-methylated catecholamines that act as tumour biomarkers (homovanillic acid and methoxytyramine^47^). We perform an in-depth characterization of different RolR variant attributes such as response, crosstalk, dynamic range, and leakiness, followed by transplantation of the engineered aTFs from *E. coli* to the model eukaryotic system *Saccharomyces cerevisiae*. This work provides a framework to perform a gradual walk across the fitness landscape of the RolR scaffold in a high-throughput manner to generate variants with altered ligand-binding properties that could function as biosensors towards sensing molecules previously undetectable by the wild-type (WT), and is broadly applicable to other aTFs and ligand-binding proteins.

**Fig. 1.**
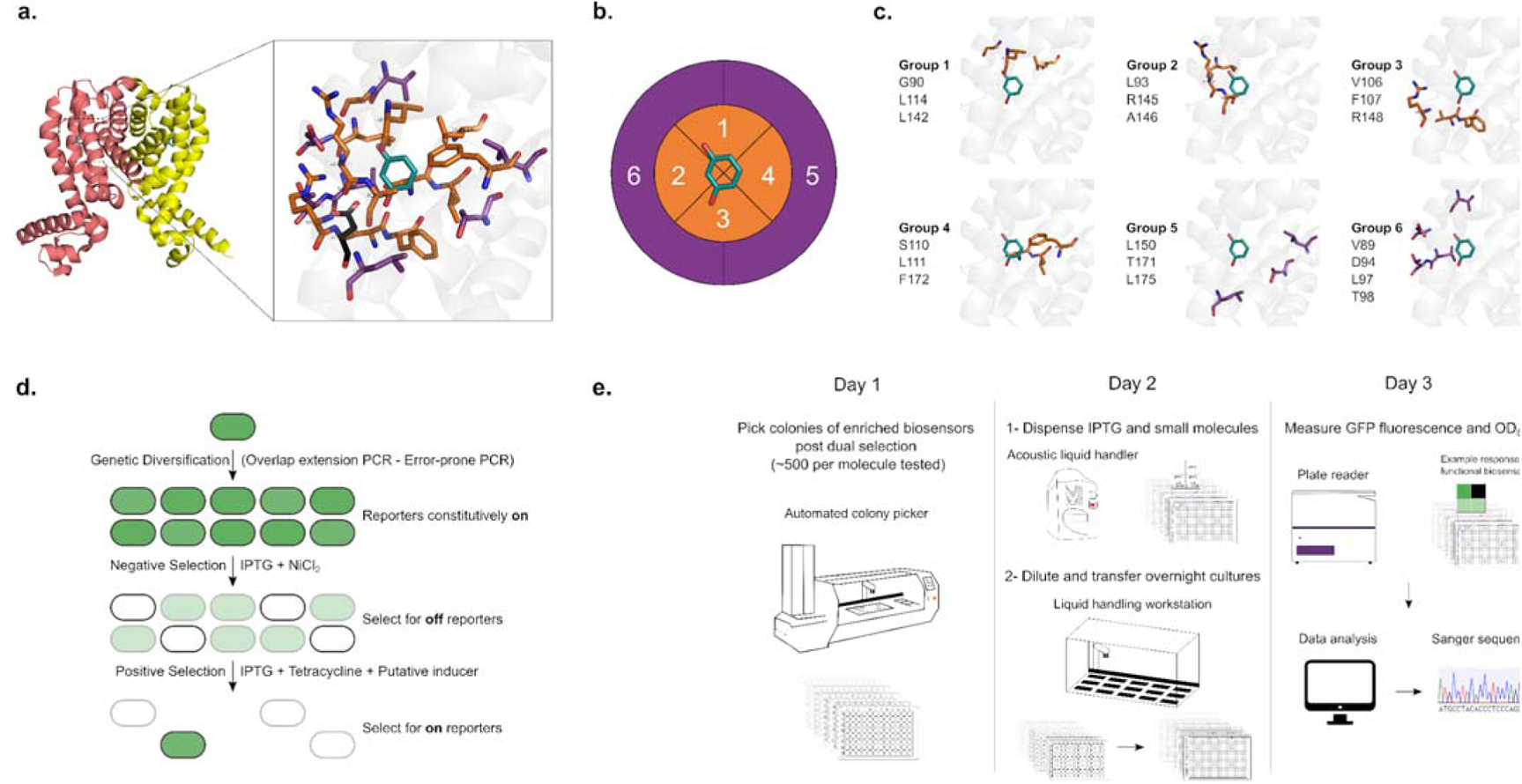
RolR mutagenesis, selection, and semi-automated high-throughput screening workflows. **a.** RolR dimer in the holo conformation (PDB:3AQT), and the structure of the ligand-binding pocket with residue D149 in black, resorcinol in teal, and the19 residues chosen for mutagenesis within 5Å in orange, and those between 5Å and 8Å in purple. **b.** Cartesian binding pocket map for combinatorial active site saturation test (CAST). **c.** The six amino acid groups comprising the 19 residues to be targeted for mutagenesis. **d.** Principle of biosensor TetA dual selection using NiCl_2_ for negative selection towards transcriptional repression competency, and tetracycline for positive selection towards the induction by ligands of interest. **e.** Semi-automated high-throughput screen. On day 1, ~500 colonies were picked for each candidate molecule. On day 2, IPTG and small molecules were dispensed using an acoustic liquid handler into 384 well plates. Growing colonies were diluted and dispensed into 384 plates testing the different states of the sensor using a liquid handling workstation. On day 3, fluorescence was measured using a plate reader. Data was then analyzed, and positive clones were genotyped by Sanger sequencing.

## Results

### Construction and inducer-characterization of resorcinol biosensors.l

RolR acts by negatively regulating the *rolHMD* cluster involved in resorcinol catabolism in *C. glutamicum*^33,34^. We have introduced the *rolR*-*rolO* repressor-operator pair into a previously described genetic circuit^48^, whereby a regulatory module comprising *rolR* is expressed from the IPTG-inducible *tac* promoter, and a reporter module is expressed from a second, RolR-regulated promoter (Fig. 2a). The engineered circuit comprises the modified promoter *P_2_*, which contains the consensus promoter elements (−35: TTGACA, −10: TATAAT) and 38 bp that include the *rolO* operator element immediately downstream of the −10 sequence (Fig. 2a). This circuit design allows the reporter module to be expressed in the absence of regulator binding, and to be repressed when RolR is produced upon induction of the regulatory module by IPTG supplementation, as seen in the gradual decrease in GFP fluorescence on increasing IPTG concentration (Fig. 2b). Adding the RolR inducer resorcinol, in the presence of IPTG, causes the gradual induction of the reporter module and increasing GFP fluorescence. This circuit setup enables tuning of EC_50_ values for resorcinol from the mid nM to low μM range (Fig. 2b).

**Fig. 2.**
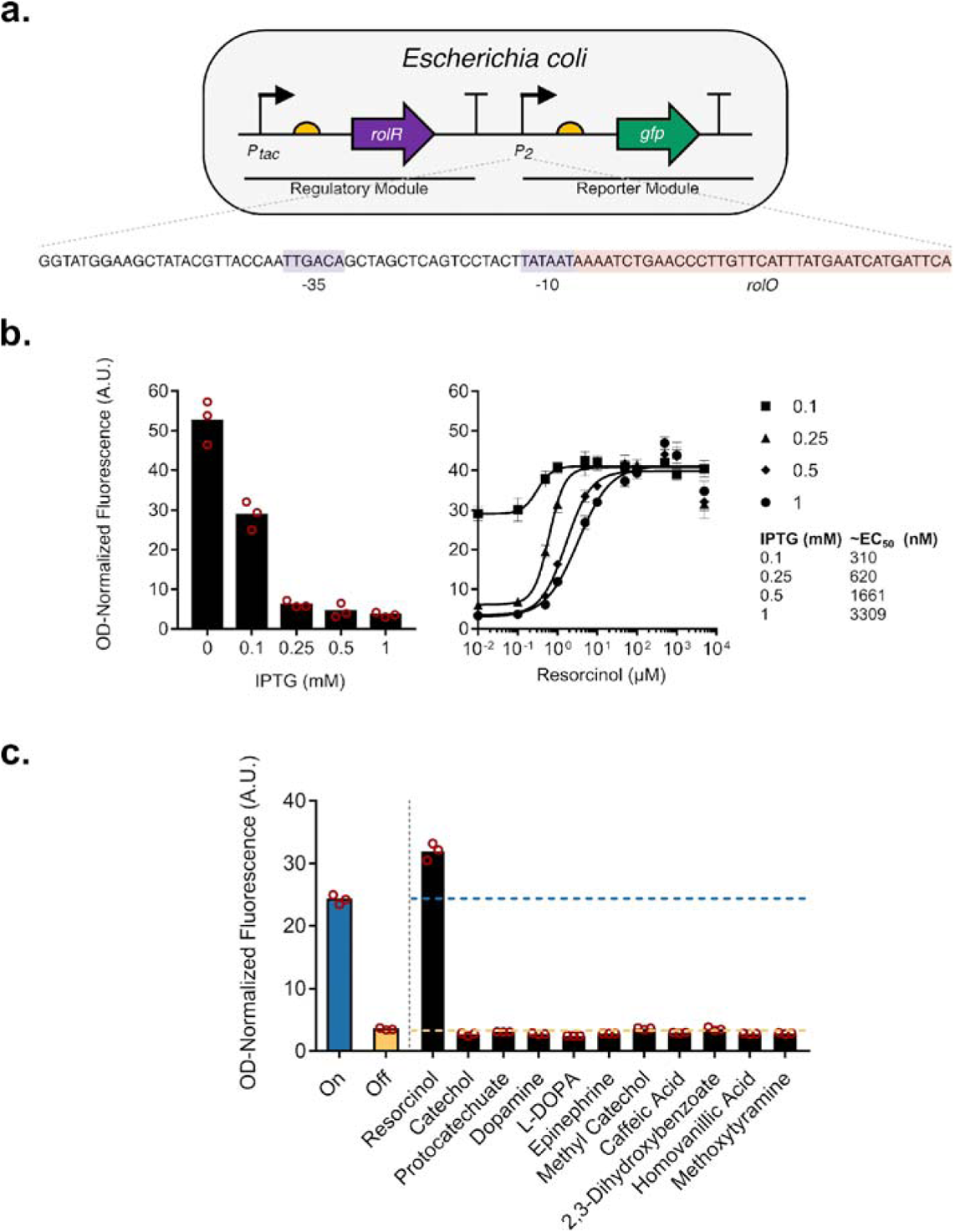
Description and functional characterization of RolR resorcinol biosensors in *Escherichia coli.* **a.** Structure of the one-plasmid RolR biosensor genetic circuit. RolR expression is driven by the IPTG-inducible *tac* promoter (regulatory module), and GFP expression is driven by a second engineered resorcinol-responsive promoter (reporter module), which contains one copy of the *rolO* operator sequence inserted immediately downstream of the −10 promoter element. **b.** Left panel: Gradually increasing IPTG concentrations reduces whole-cell OD-normalized fluorescence levels. Right panel: Gradually increasing resorcinol concentrations in fixed IPTG concentrations results in an increase in OD-normalized fluorescence levels, and resorcinol EC_50_ values in different IPTG concentrations were measured. **c.** Ligand-specificity of RolR towards all molecules in this study in the presence of 1 mM IPTG. Datapoints are mean ± SD, *n* = 3. A.U., arbitrary units.

Most aTFs have evolved towards specific ligands, yet some exhibit promiscuity towards multiple small molecules, and are sometimes called “generalists”^24,49–51^. Thus, we sought to determine if expression from *P_2_* could be induced by other molecules of interest. We have tested the RolR circuit by introducing a range of aromatic small molecules containing catechol moieties and did not observe induction by any molecules other than resorcinol (Fig. 2c).

### A selection-screening framework to identify RolR variants that respond to new ligands

Many successful aTF directed evolution efforts have resulted in expanding ligand specificity towards new molecules^52–54^. This motivated us to create high quality variant libraries that sample the protein sequence landscape more effectively, in an attempt to perform a gradual walk across the fitness landscape towards molecules with divergent structures (see “Methods”). Some amino acid substitutions in aTFs can affect their DNA binding affinity, resulting in altered transcriptional regulation^52,55^. Dual selection schemes are essential for aTF directed evolution to select for and maintain DNA-binding competency (transcriptional repression), and induction by the ligand of interest, the former which could be easily perturbed by substitutions due to the nature of allostery^31^. Fluorescence activated cell-sorting has been used extensively for dual selection^53,56^, and genes that encode multiple phenotypes such as TolC^57^ and thymidine kinases^58^ have been used as dual selection markers. To this end, we employ the *E. coli* TetA antiporter as a dual selection marker to couple aTF activity to cell growth^38,39,59^, which requires tetracycline and NiCl_2_ for positive and negative selection, respectively^60^. We extended the reporter module by inserting TetA downstream of GFP for polycistronic expression (Fig. 3a), and characterized the system in both liquid and solid media. In this selection scheme, negative selection is first employed to select for DNA-binding-competent variants of RolR that repress TetA expression, allowing the cells to survive in media supplemented with NiCl_2_^61^. This is then followed by positive selection to identify variants that are induced by a ligand of interest, where induction of TetA expression causes a tetracycline resistant phenotype (Fig. 3b). In a liquid medium, we identified the concentrations for selection with the most growth difference between repressed and induced states to be 0.4 mM NiCl_2_ (Fig. 3c, d) and 7.5 µg ml^−1^ or higher for tetracycline (Fig. 3e, f). As well, given that TetA and GFP are expressed from the same promoter, we observed a correlation between the expected NiCl_2_ and tetracycline resistance or susceptibility phenotypes and cell population fluorescence levels (Fig. 3g). Furthermore, on solid medium, negative selection performed better at a higher concentration of NiCl_2_ (0.6 mM) compared to liquid medium, whereas positive selection on tetracycline performed equally on liquid and solid medium (Fig. 3h, i).

**Fig. 3.**
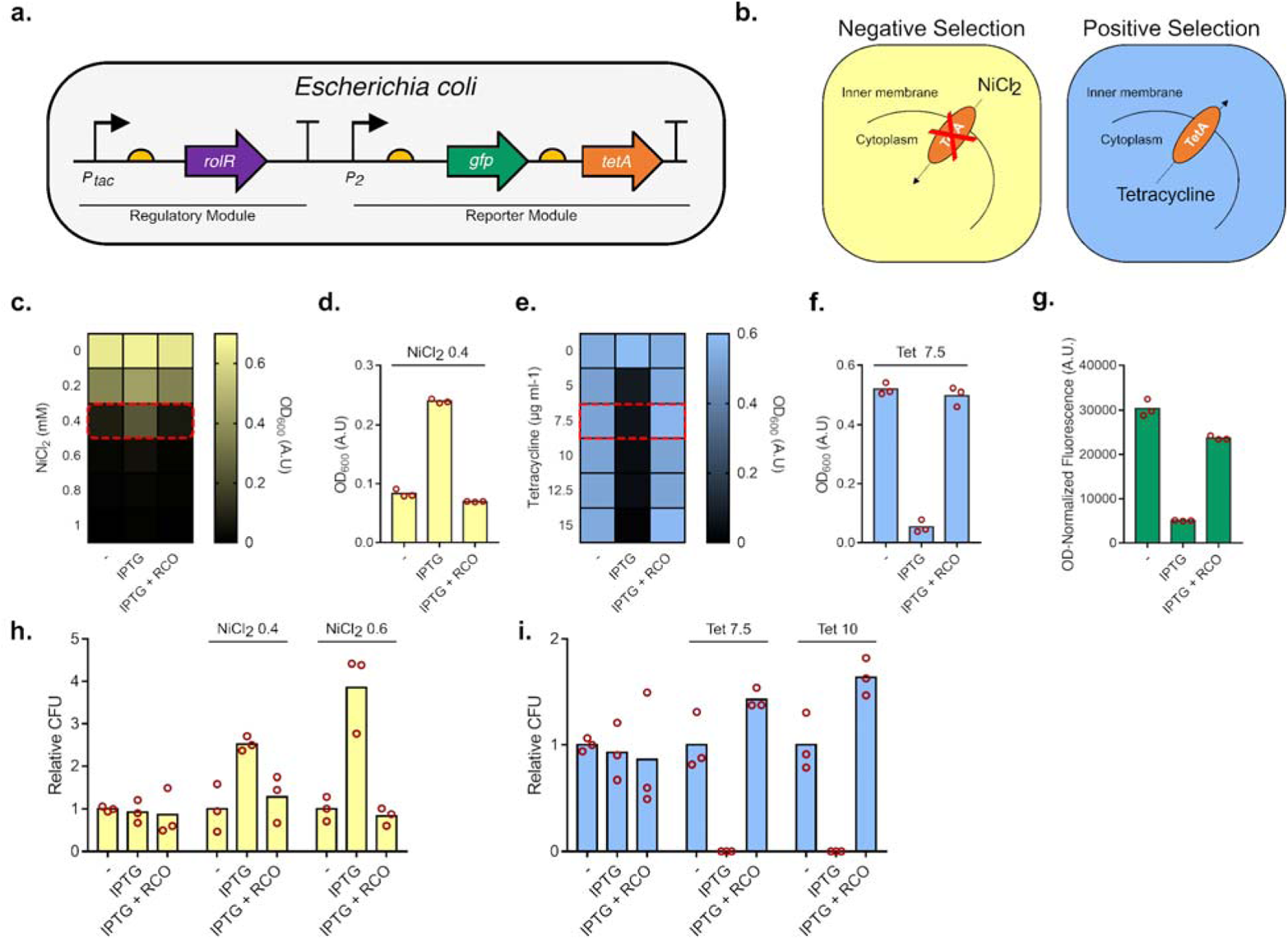
Characterization of the TetA dual selection system. **a.** The reporter module was extended by introducing TetA downstream of GFP. **b.** Mechanism of TetA dual selection. Negative selection selects for the ability to repress expression from promoter *P_2_* which would not allow for toxic NiCl_2_ to enter the cell. Positive selection selects for induction of expression from *P_2_*by a ligand of interest, which would allow for tetracycline resistance. **c.** Characterization of negative selection in liquid media in different NiCl_2_ concentrations, and **d.** in 0.4 mM NiCl_2_ (RCO=resorcinol). **e.** Characterization of positive selection in liquid media at different tetracycline concentrations, and **f.** in 7.5 µg ml^−1^ tetracycline. **g.** GFP fluorescence levels correlate with expected phenotypes associated with TetA expression. **h.** Negative selection in solid media in 0.4 and 0.6 mM NiCl_2_. **i.** Positive selection in solid media in 7.5 and 10 µg ml^−1^ tetracycline. Datapoints are mean ± SD, *n* = 3. A.U., arbitrary units. CFU, colony forming units.

### First generation variants towards catechol

In the first generation of variant library hits, we identified 19 unique variants that are induced by catechol to varying degrees (Fig. 4a, Supplementary Fig. 1a). Variants induced by catechol show an overrepresentation of substitutions in amino acid positions within groups 1 and 6, which are spatially close to the OH groups of resorcinol (Fig. 4a, Supplementary Fig. 1a). Our top-performing catechol-responsive variants, named CAQ101, CAQ102, CAQ.Comb.4, and CAQ.Comb.5, had maximum induction values of ~4-5 fold in catechol, while retaining responsiveness to resorcinol (Fig. 4b, Supplementary Fig. 1b). We found variant CAQ101 to have a bell-shaped response, seen in the increase in fluorescence followed by a subsequent decrease in increasing catechol concentrations. This type of response was previously described, yet the precise mechanism behind it remains unknown^52,62^. Variants from this library also exhibited varying dynamic ranges defined here as the difference in signal between on and off states in the absence and presence of IPTG, respectively (Fig. 4c, Supplementary Fig. 1d). We have also observed that all 19 identified variants with increased response to catechol always have substitutions in at least one of positions V89, L93, or L142, with substitutions to the aromatic hydrophobic residues phenylalanine, tyrosine, or in only one case to hydrophobic aliphatic isoleucine. These residues cluster close to the top OH group of resorcinol, which would suggest that the substitutions contribute by slightly reshaping the pocket to accommodate the hydroxyl group of a benzenediol isomer of resorcinol at an *ortho* position relative to the bottom OH group, and as a result being able to accommodate catechol. Furthermore, when combining these substitutions, only variants CAQ.Comb.4 and CAQ.Comb.5 had a significant induction fold change in their response to catechol, as well as a functional dynamic range (Fig. 4b, c, Supplementary Fig. 1c).

**Fig. 4.**
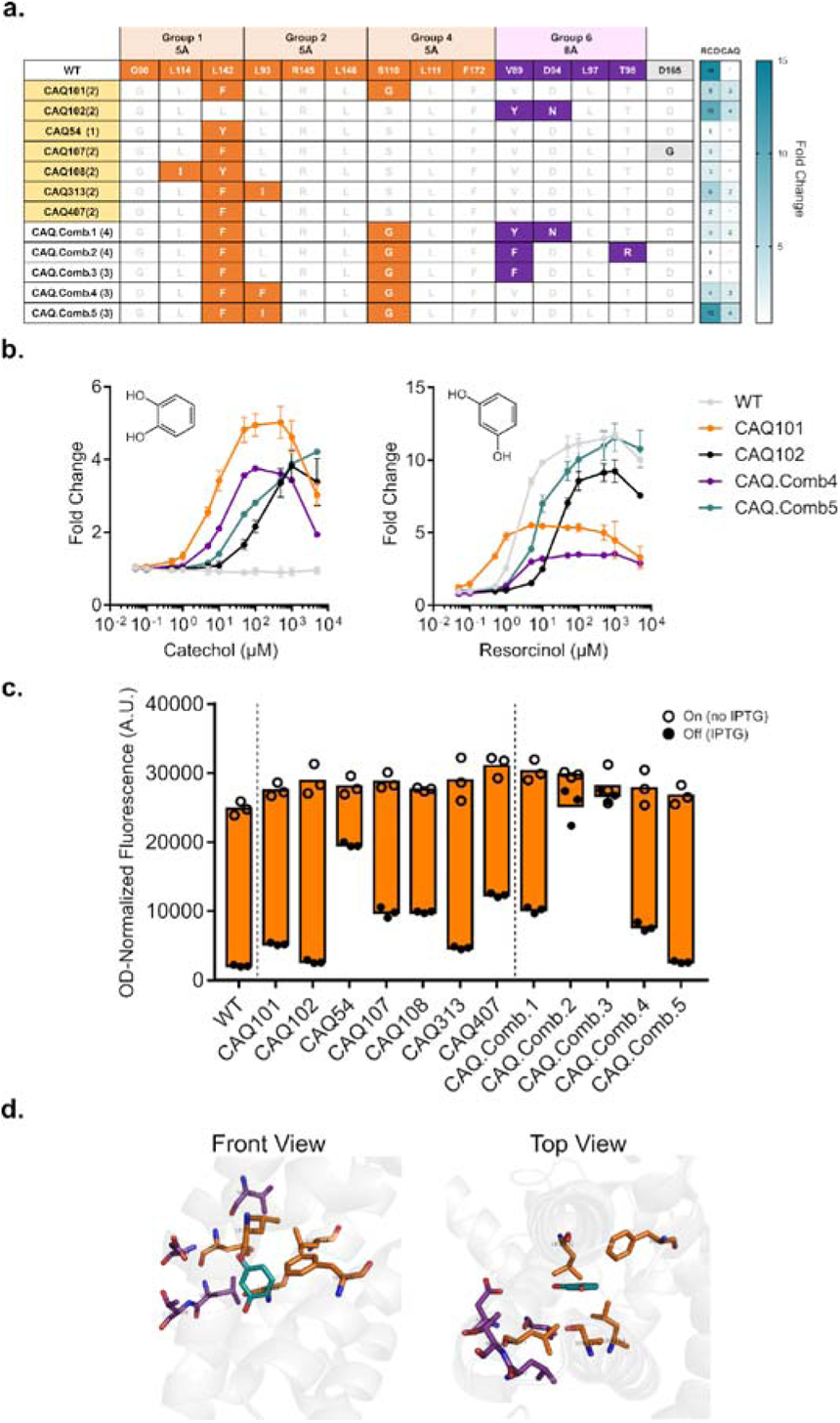
First generation variant hits towards catechol. **a.** Table depicting a subset of identified variants responsive to catechol, their substitutions coloured according to group, and their response towards resorcinol (RCO) and catechol (CAQ). Variants highlighted in yellow were used as a template for further mutagenesis. **b.** Dose response curves of the top-performing variants compared to WT RolR with catechol (left panel) and resorcinol (right panel). **c.** Dynamic range of WT RolR and identified variants. **d.** Position of residues mutated in the RolR binding pocket around resorcinol using PDB:3AQT. Datapoints are mean ± SD, *n* = 3. A.U., arbitrary units.

CAQ.Comb.5 does not exhibit the bell-shaped response that is seen in its parent variant CAQ101, and the loss of that effect was achieved by the single additional substitution L93I (Fig. 4a-c), unlike CAQ.Comb.4, which contains a phenylalanine residue at position 93 and retains the bell-shaped response. Lastly, some of the variants were found to have a D165G substitution, which was due to a gene synthesis error that appeared to slightly reduce leakiness (off state), which was observed when comparing the responses of variants CAQ107 and CAQ407 (Fig. 4a, c).

### Second and third generation variants towards substituted catechol moieties

In addition to catechol, we sought to add diversity to the range of recognized inducer molecules by identifying variants that respond to catechol derivatives with substitutions in positions 3 and 4 of the benzene ring. For the second generation library, we used 7 variants from the first library as templates (Fig. 4a). We then created variant libraries targeting groups 3, 4, and 5, which align with the hydrophobic side of the ring where the required substitutions occur. Furthermore, we revisited the structure of RolR and identified the additional and adjacent residues of interest L142, G143, and A146, as well as targeted a modified group 1 that included residues G90, L114, and F172. Following dual selection and screening, we identified variants that were enriched from pools where methyl catechol, caffeic acid, or L-DOPA were used for positive selection, and that displayed activity towards these molecules. We also identified variants enriched in the epinephrine-, dopamine-, and protocatechuate-selected pools that displayed no change in GFP expression in the presence of these molecules (Supplementary Fig. 2). We used a subset of these variants, in addition to the catechol-responsive variant CAQ101, as a template for a final round of random mutagenesis and selection targeting the entire ligand-binding domain of RolR, in a strategy similar to that used to improve the response of weak biosensors (Supplementary table 2)^55^. We then characterized the identified variants in terms of response, crosstalk, dynamic range, and we also investigated the position of substitutions in the binding pocket.

For variants selected for response to catechol, the third generation variant CAQ3 notably lacked the bell-shaped response of CAQ101 (Fig. 5a). Furthermore, CAQ3 behaved favourably by having significantly reduced the crosstalk with methyl catechol and caffeic acid that was seen with CAQ101, as well as by having reduced leakiness (Fig. 5a). This was all achieved by a single distal M139L substitution, wherein methionine is replaced with another hydrophobic residue, leucine (Fig. 5a, Supplementary table 2). In the case of methyl catechol, the second and third generation variants MC2 and MC3 had a similar response, although it was not possible to test higher concentrations of methyl catechol due to its toxicity to *E. coli* (Fig. 5b). Both variants exhibited minimal crosstalk with other molecules, particularly catechol, but MC3 had a significantly improved dynamic range, which is a desirable attribute for aTF-based biosensors (Fig. 5b). This was achieved also by the single substitution H113L, rendering this position hydrophobic (Fig. 5b, Supplementary table 2). For L-DOPA, the second and third generation variants LD2 and LD3.1, respectively, both exhibited ~2 fold increase in fluorescence in the presence of L-DOPA, with limited crosstalk, near wild-type dynamic range levels, and with most substitutions clustering around the hydrophobic side of the ring (Fig. 5c, Supplementary table 2). In the case of the tumour marker homovanillic acid, which differs from the rest of the molecules in that the phenol has an *ortho*-methoxy substitution in addition to a *para*-carboxymethyl substitution, we observed a minimal bell-shaped response that reached a maximum induction of ~1.5 fold at 0.1 mM homovanillic acid, as well as minimal crosstalk and a dynamic range that is approximately that of the wild-type (Fig. 5d). Substitutions in variant HVA3 to accommodate this molecule were mainly in the newly substituted adjacent residues 142, 143 and 146 (Fig. 5d, Supplementary table 2). For caffeic acid, the second generation variant CA2 performed poorly, with better variants identified in the third generation (Fig. 5e). All caffeic acid-responsive variants identified in the third generation exhibited significant crosstalk with catechol and less so with methyl catechol, except for variant CA3.3, which, as a tradeoff, exhibited a poorer dynamic range relative to other third generation variants (Fig. 5e). Substitutions were once again found to cluster around the hydrophobic side of the ring where the substitution occurs (Fig. 5e, Supplementary table 2). Lastly, of all identified aTF variants, the protocatechuate-responsive variant PCA3 appeared to be the weakest of all, as GFP expression is induced by approximately 1.2 fold, which is further weakened by its bell-shaped response (Fig. 5f). PCA3 exhibited limited crosstalk, an acceptable dynamic range, with substitutions clustering around the hydrophobic side (Fig. 5f), and with the same substitutions seen in variant LD3.2 (Supplementary table 2). Lastly, we identified variants that exhibited an abolished response to the cognate inducer, resorcinol, such as LD2 (Supplementary Fig. 3, 4).

**Fig. 5.**
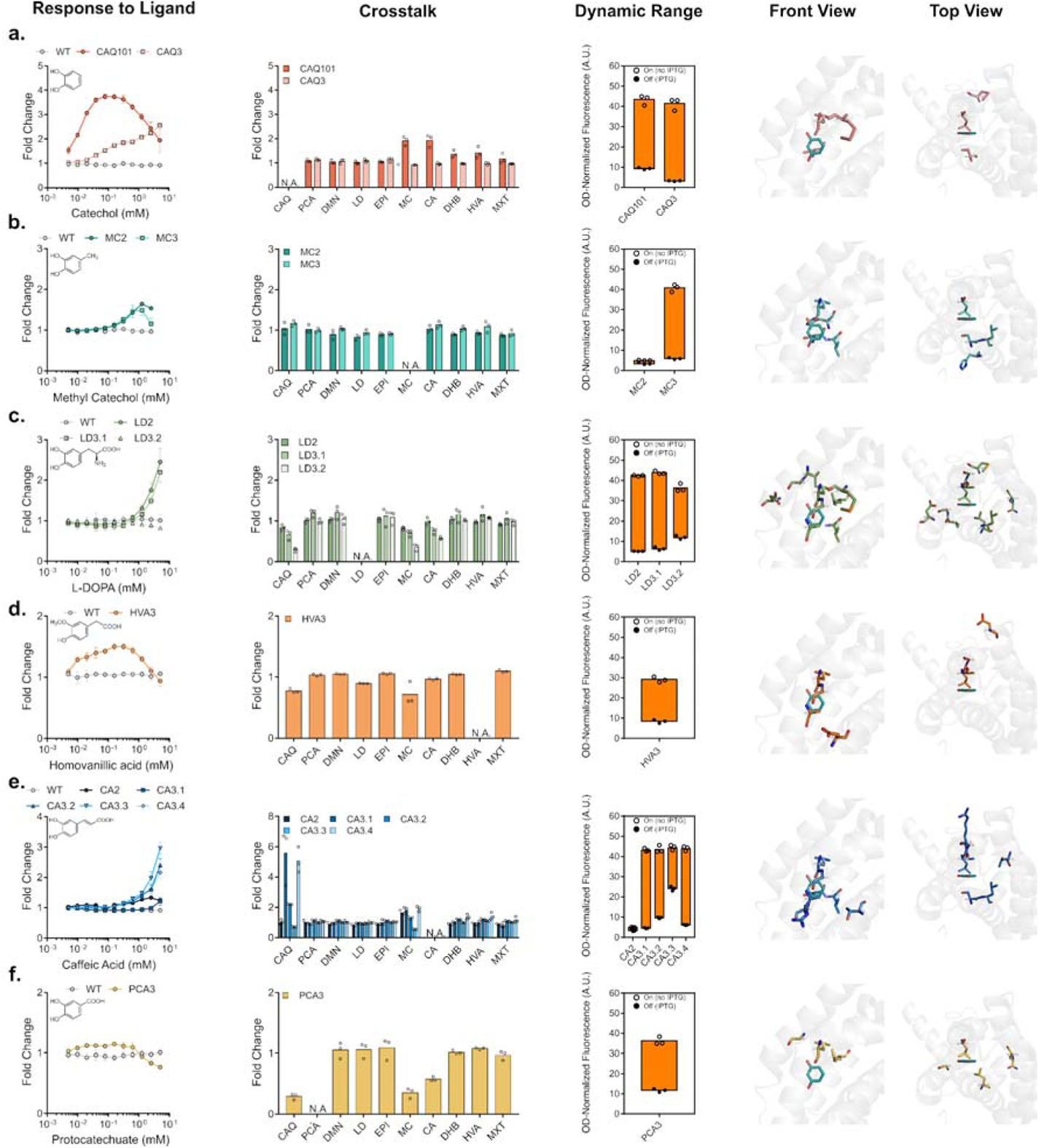
Second and third generation variant hits. Characterizing variant biosensor attributes of response to ligand, crosstalk (CAQ=catechol, PCA=protocatechuate, DMN=dopamine, LD=L-DOPA, EPI=epinephrine, MC=methyl catechol, CA=caffeic acid, DHB=2,3-dihydroxybenzoate, HVA=homovanillic acid, MXT=methoxytyramine), dynamic range, and the positions of the identified substitutions in the binding pocket, respectively, for variants identified for catechol **(a)**, methyl catechol **(b)**, L-DOPA **(c)**, homovanillic acid **(d)**, caffeic acid **(e)**, and protocatechuate **(f)**. Datapoints are mean ± SD, *n* = 3. A.U., arbitrary units.

### Transplantation of biosensor in *Saccharomyces cerevisiae*

Bacterial aTFs have been successfully transplanted into eukaryotes, such as the yeasts *S. cerevisiae* and *Yarrowia lipolytica*, in addition to mammalian cell lines^63–68^. Here, to demonstrate the portability of RolR and its variants, we transplanted a subset of RolR variants into genetic circuits in *S. cerevisiae*. We integrated a single copy of our genetic circuit, which consists of a regulatory module comprising RolR expressed from the strong constitutive yeast promoter *P_TDH3_* (Fig. 6a). For the reporter module, we employed the *P_CCW12_* promoter to drive expression of the Envy fluorescent protein^69^. We engineered the promoter for RolR regulation by inserting a single copy of the *rolO* operator 11 bp downstream of the terminal consensus TATA box sequence of the promoter, TATAWAWR (W=A/T R=A/G), as previously described^70^. The WT RolR biosensor was functional in *S. cerevisiae*, in which we identified additional methods for tuning the response curve of genetic circuits, namely codon optimization and the type of media used. This gave rise to a maximal induction of ~8 fold change in the minimal YNB medium with the codon-optimized RolR variant (Fig. 6b). As seen when using *E. coli* as host, WT RolR is specific to resorcinol in the *S. cerevisiae* system, with no observable response towards any of the other molecules in this work at concentrations up to 5 mM (Fig. 6c). Of all our generated variants, the catechol-responsive variant CAQ101 exhibited the strongest response to its cognate inducer (~10 fold change), but without the bell-shaped response phenotype that was seen in *E. coli* (Fig. 6d).

**Fig. 6.**
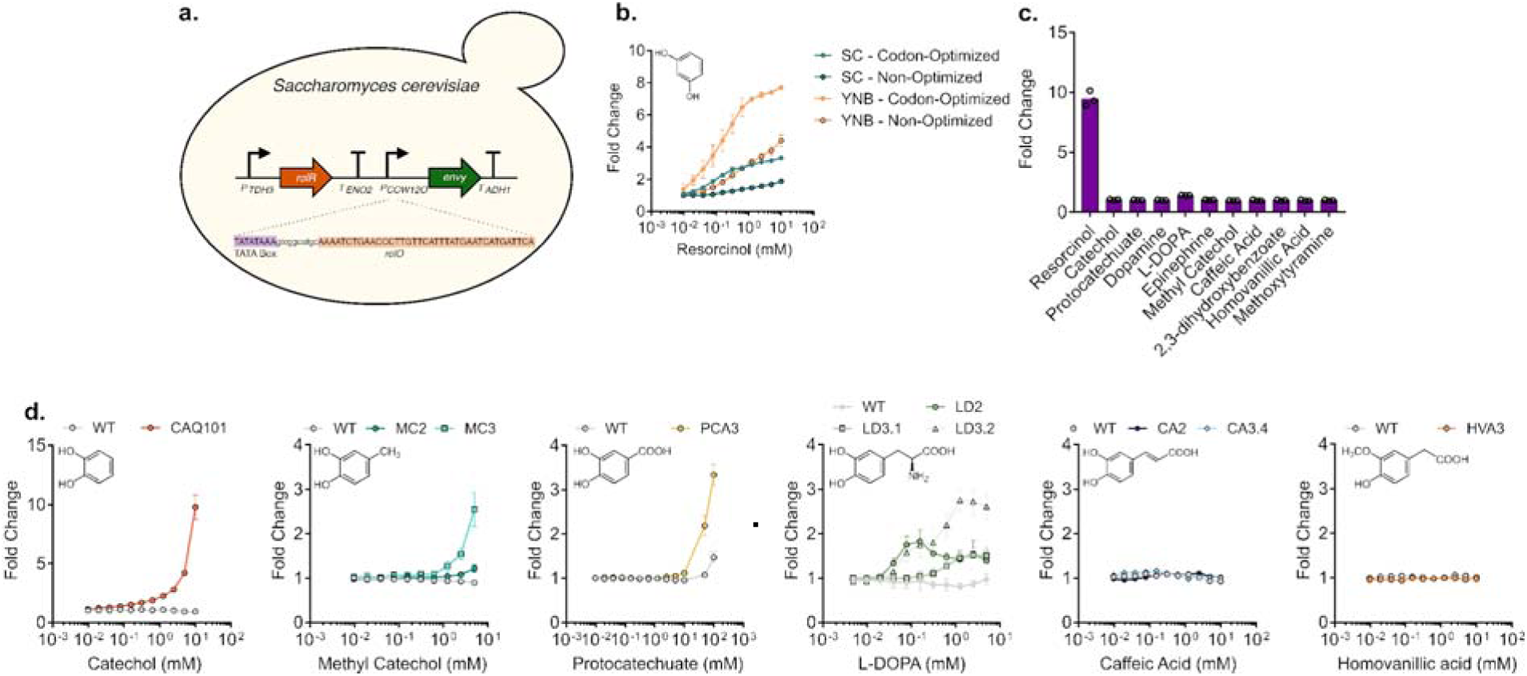
Construction and characterization of wild-type and variant RolR biosensors in *Saccharomyces cerevisiae.* **a.** Design of chromosomally integrated RolR genetic circuit in S. *cerevisiae*. RolR is expressed from the strong constitutive promoter *P_TDH3_*, and Envy GFP is expressed from the *P_CCW12_* promoter, which was engineered for resorcinol-responsiveness by inserting 1 copy of *rolO* downstream of the TATA box. **b.** Response of the resorcinol biosensor in *S. cerevisiae* and tuning of the circuit response by altering codon-optimization status and inoculation media. The sensor exhibited the best induction and EC_50_ when codon-optimized for yeast expression and when minimal YNB media was used. **c.** Response of the WT RolR with all the small molecules included in this study. **d.** Dose-response curves of representative variant RolR biosensors compared to WT RolR with catechol, methyl catechol, protocatechuate, L-DOPA, caffeic acid, and homovanillic acid. Datapoints are mean ± SD, *n* = 3.

Methyl catechol-responsive variants MC2 and MC3 are both functional, with the ~3 fold induction of the third generation variant MC3 significantly better than the response of MC2 (Fig. 6d). PCA3 exhibited a modest 1.2 fold change at 10 mM protocatechuate but when the concentration was increased to 100 mM, which is possible due to the tolerance of *S. cerevisiae* to this molecule, a ~3 fold induction was observed relative to ~1.5 fold from the WT, and, similar to CAQ101 without the bell-shaped response observed in *E. coli* (Fig. 6d). Furthermore, L-DOPA-responsive variants also exhibited a ~1.5-3 fold induction in 5 mM L-DOPA (Fig. 6d). However, variant LD3.2, which exhibited the strongest response in *S. cerevisiae* to L-DOPA lacked an observable response in *E. coli*, which highlights possible variations in biosensor response in different microbial chassis. Lastly, we could see no response with the generated caffeic acid- and homovanillic acid-responsive variants when transplanted to *S. cerevisiae*, further emphasizing these differences (Fig. 6d).

It has been shown that introducing substitutions in aTF repressors can reverse the activities towards their ligands, causing them to exhibit a stronger affinity to DNA in the presence of their ligand instead of the loss of affinity to DNA that occurs under normal induction conditions. This was seen with the introduction of 2 substitutions in TetR to generate revTetR, which is used in the Tet-OFF system for regulating gene expression in response to tetracycline and its analogues^71^. Furthermore, some aTFs exhibit a dual function with different molecules such as TyrR, which has its regulated promoter activated by phenylalanine and repressed by tyrosine^72^. Here, on characterizing variant PCA3 in *E. coli*, in addition to the weak, 1.2-fold bell-shaped induction response to protocatechuate, we noticed a significant reduction in fluorescence in response to catechol (Figure 5, Supplementary Fig. 5a). When the PCA3 circuit was introduced to *S. cerevisiae*, we observed a robust and stronger response, where we identified what strongly suggests three stable and distinct expression states from that promoter: an intermediate state in the absence of any effector molecule, an induced state in presence of protocatechuate, and a repressed state in the presence of catechol (Supplementary Fig. 5b). Titrating these molecules yielded a significant response in *S. cerevisiae* to both molecules, but not in *E. coli* (Supplementary Fig. 5c, d).

## Discussion

In the present work, we engineered a diverse collection of aTF-based biosensors with expanded specificity and minimal crosstalk towards aromatic molecules by combining a structure-guided approach with a dual selection scheme and automated screening. We generated and screened small variant libraries that sampled a fraction of the aTF’s sequence space with mutations focused around the resorcinol-binding region of RolR followed by randomized libraries targeting the ligand-binding domain. This protein engineering/evolution approach resulted in biosensors that respond to catechol, protocatechuate, and caffeic acid, in addition to the molecules homovanillic acid, L-DOPA, and methyl catechol for which whole-cell aTF biosensors had not been reported to date.

Directed evolution campaigns to evolve generalist aTFs to be more specific towards a single target molecule show a high rate of success^50,56,73^. Evolving narrow or ligand-specific aTFs for induction by new molecules was also recently reported using one or a combination of random mutagenesis, computational protein design, and site saturation mutagenesis^52,53, 74–76^. Similar to other proteins, evolving new activities in ligand-specific aTFs is traditionally more challenging, and is facilitated by both structural knowledge and chemical intuition^77,78^. Here, we demonstrate the evolvability of the resorcinol-specific aTF RolR (Fig. 2c) by engineering biosensor variants that respond to six molecules that are not recognized by the parent protein and with minimal crosstalk between ligands (Fig 5). This demonstrates the benefit of structure-guided site-saturation mutagenesis of the ligand-binding pocket coupled with random mutagenesis of the entire ligand-binding domain to simultaneously evolve a single ligand-specific aTF to respond to multiple molecules. Similar methods were used for example to improve response and eliminate crosstalk of the promiscuous aTFs RamR^73^ and LysG^56^, or to expand substrate specificity of AraC^74^,PcaV^75^,and VanR^76^ and over a single generation of mutagenesis.

Of the six molecules RolR variants were evolved to respond to in this study, five had variants that were identified intentionally (*e.g.* a catechol responsive variant from the catechol-selected pool) and one molecule, protocatechuate, had the variant PCA3/LD3.2 identified incidentally as responsive to a molecule that was not used for its selection. Variant PCA3/LD3.2 responds to both protocatechuate and L-DOPA, but was identified from a pool that underwent selection using methoxytyramine. This particular variant emphasizes a general issue with aTF engineering that could be regarded as a double-edged sword. PCA3/LD3.2 likely escaped our dual selection due to its leaky off state (or intermediate state considering its tri-stability) (Fig. 5f), which would have positioned this state in a sweet spot that is able to survive both negative and positive selection without entirely repressing transcription nor being induced by the inducer of interest. As a comparison, this variant would have been missed in the extensively used fluorescence-activated cell sorting methodologies, where the highest and lowest signallers are usually selected for, with intermediate variants entirely eliminated^53,56^. This overlooks the inherent complexity of allostery, and leads to losing significant diversity from the variant library that could be seen in leaky variants as well as intermediate signallers. This bifunctional/tri-stable biosensor PCA3/LD3.2 was identified as an unintentional outcome of our selection method, which, in an effort to eliminate crosstalk was modified in the third generation to include a negative selection step including all molecules from our panel other than our desired target for evolution. Adding catechol to our negative selection pools led to the identification of this biosensor that repressed expression in the presence of catechol, and induced it in the presence of protocatechuate, with an intermediate state of promoter expression in the absence of either molecule. These findings suggest a new and tantalizing methodology for aTF engineering, wherein instead of evolving aTFs for activity towards a single molecule, one could evolve them directly for induction by one molecule and repression by another. This would ultimately add to the *in vivo* pathway dynamic control biosensor toolbox in the field of metabolic engineering^79^, and, more importantly, is truly inspired by nature, which has already evolved bi-functional aTFs^72^.

We were able to observe a response of our evolved RolR towards the tumour marker homovanillic acid (Fig. 5d). This molecule along with methoxytyramine could be used for screening of catecholamine-secreting tumours such as paraganglioma, phaeochromocytoma, among others, as well as for monitoring disease progression following treatment^47^. We observed maximum fluorescence induction at 0.1 mM homovanillic acid, or ~18 mg L^−1^, which is higher than the limit of detection of current diagnostic methods (0.03 - 0.3 mg L^−1^)^80^. Additional protein engineering or the use of cascaded amplification circuits^21^ will be needed to enhance the sensitivity and response to homovanillic acid to enable clinical applications of this sensor^80,81^.

Transplanting aTFs into *S. cerevisiae* has been accomplished using repressors^63^, aporepressors^82^, and activators^64^. Engineering promoters for regulation by repressors and aporepressors have the advantage of being more straightforward than activators, as they require the operator to be placed downstream of the terminal TATA box of the promoter, compared to a variable and unique location for every activator^70,82,83^. Here, to demonstrate the versatility of RolR and the generated variants, we transplanted them as components of engineered genetic circuits into *S. cerevisiae*, and observed responses that were generally stronger than those observed from *E. coli* (Fig. 6). This observation could be a result of differences in small molecule membrane transport between these organisms or differences between the biosensor contexts in both organisms, which exist as a multi-copy plasmid in *E. coli* and a single-copy genomic integration in *S. cerevisiae.* This was observed specifically with variant LD3.2, which exhibited undetectable activity for L-DOPA in *E. coli* but was functional in *S. cerevisiae*, as well as with the tri-stable switch PCA3. Conversely, sensors for homovanillic acid and caffeic acid showed no activity in *S. cerevisiae* compared to *E. coli*. In particular, we have previously seen caffeic acid-responsive biosensors in *E. coli* that did not function in *S. cerevisiae*, further emphasizing these differences^24^.

We were not able to engineer RolR to respond to the catecholamines dopamine, epinephrine, and the tumour marker methoxytyramine. The only catecholamine that gave a response was L-DOPA, which has a terminal carboxylic acid and an internal amine group, as opposed to the terminal amine group seen in the other targeted catecholamines. We hypothesize that this terminal amine could be the contributor behind the failure to identify variants that respond to them, and that further structure-guided engineering will be needed to engineer RolR response to catecholamines. We also were not able to identify variants responsive to 2,3-dihydoxybenzoate, which is the only benzene derivative in this work with substitutions on positions 1, 2, and 3, as opposed to 1-, 2-, and 4-substitutions for the other trisubstituted benzenes, suggesting that different residues will have to be targeted to evolve response towards substitutions in these positions.

Our methodical approach for engineering aTFs provides a process for iteratively and systemically traversing their fitness landscapes to acquire novel ligand specificities. This approach has the potential for dramatically increasing the repertoire of inducer molecules for aTFs that are functional in broad host backgrounds, subsequently expanding opportunities for small molecule-directed programming of cellular activity in many metabolic engineering, cell, and synthetic biology applications. Additional directions for RolR engineering could focus on expanding its biosensor capabilities towards further diverse structures, such as the resorcinol-containing stilbene moiety seen in cannabinoid precursors^84^, or the catechol-containing tetrahydroisoquinoline alkaloids^45^, as well as other tumour biomarkers such as metanephrines.

## Methods

### Strains and growth media

*E. coli* DH5α was used for all molecular cloning, plasmid propagation, genetic circuit characterization, and protein engineering and directed evolution experiments. Unless otherwise stated, *E. coli* cells were cultured at 37 °C for ~16 h in liquid or solid LB medium (10 g L^−1^ NaCl, 10 g L^−1^ tryptone, 5 g L^−1^ yeast extract). Biosensor selection and response characterization was performed in M9 minimal medium (6.78 g L^−1^ Na_2_HPO_4_, 3 g L^−1^ KH_2_PO_4_, 1 g L^−1^ NH_4_Cl, 0.5 g L^−1^ NaCl, 4 g L^−1^ Dextrose, 2 mM MgSO_4_, and 0.1 mM CaCl_2_) supplemented with 0.34 g L^−1^ thiamine hydrochloride, 2 g L^−1^ casamino acids, and 100 mM ascorbic acid as an antioxidant. Kanamycin (50 μg ml^−1^) was added for plasmid selection as needed.

*S. cerevisiae* manipulations were conducted in the prototroph CEN.PK113-7D, which was grown at 30 °C in liquid or solid Yeast Peptone Dextrose medium (YPD; 10 g L^−1^ yeast extract, 20 g L^−1^ peptone, 20 g L^−1^ dextrose) for ~16 h or 48 h, respectively. Biosensor characterization was performed in Yeast Nitrogen Base medium (YNB; 6.7 g L^−1^ yeast nitrogen base, 20 g L^−1^ dextrose) and Synthetic Complete medium (SC; 6.7 g L^−1^ yeast nitrogen base, 20 g L^−1^ dextrose, 1.92 g L^−1^ yeast synthetic drop-out medium supplement without histidine, 0.08 g L^−1^ histidine). Hygromycin (200 μg ml^−1^) and G418 (200 μg ml^−1^) were added for plasmid selection as needed.

### Chemicals

Chemicals for biosensor testing were purchased from Sigma Aldrich and dissolved in water, unless otherwise specified, to obtain the stock concentrations of 100 mM resorcinol, 100 mM catechol, 50 mM protocatechuate, 100 mM dopamine, 5 mM L-DOPA (dissolved directly in media), 100 mM epinephrine, 100 mM methyl catechol, 100 mM caffeic acid (dissolved in 99% ethanol), 50 mM 2,3-dihydroxybenzoate, 50 mM homovanillic acid, 50 mM methoxytyramine. IPTG 1 M stock solutions were prepared by dissolving in water.

### Synthetic genes, oligonucleotides, and plasmids

The *rolR* (*cgl1157*) gene from *Corynebacterium glutamicum* was codon-optimized for yeast expression using the IDT tool and was purchased from Thermo Scientific. Oligonucleotides were purchased from Thermo Scientific and suspended in nuclease-free water to a concentration of 100 μM. Sequences of oligonucleotides used in this study are listed in Supplementary data 1. Plasmids and sequences of parts used in this work are listed in Supplementary table 1 and Supplementary data 2, respectively.

### RolR genetic circuit construction

Genetic circuits were constructed using the template plasmid pQacR-Q2 (Addgene plasmid # 74690)^48^. The *qacR* gene was replaced with the codon-optimized *rolR* gene using primer sets 1-4 by polymerase incomplete primer extension (PIPE) cloning^85^ by mixing equal volumes of PCR products of *rolR* and the vector backbone each amplified without a final extension step, and each containing a 20 bp homology arm to the other part. Promoter *P_QacR_* was modified by replacing *qacO* with *rolO*, and optimizing its −10 sequence to the consensus TATAAT, creating promoter “*P_2_*”. As well, *eyfp* was mutated to *egfp*. Finally, a second ribosomal binding site followed by TetA (amplified from pBBR1MCS3) were inserted downstream of eGFP.

### RolR variant library construction

Our variant library construction strategy exploits the combinatorial active-site saturation test (CAST)^35^ and iterative saturation mutagenesis (ISM)^36^ methodologies, combined with library under-sampling^37^ in order to build focused RolR variant libraries. For mutagenesis, we selected all residues within 5Å of the inducer molecule in addition to some residues between 5Å and 8Å that we posited to have an influence on binding pocket structure as well as interactions with the inducer molecule, for a total of 19 amino acids. We chose not to mutate residue D149, as it has been established that it is essential for RolR ligand recognition through direct interaction with one of the hydroxyl groups of resorcinol, and mutating it was shown to abolish induction^34^ (Fig. 2a). We then used site saturation mutagenesis to mutate the selected positions in the ligand-binding pocket by splitting them into six groups of spatially proximal residues, which were built as separate variant libraries targeting 3-4 amino acids each (Fig. 2b, c). Then we subjected these libraries to dual selection (Fig, 2d), followed by an automated high-throughput screen to interrogate the post-selection enriched variants (Fig. 2e).

Amino acid-randomized libraries of RolR were generated using overlap extension PCR^86^. The residues of interest were randomized by inserting the NNK degenerate codon using the primer sets listed in Supplementary data 1. Overlap extension PCR products were gel-purified, PCR-amplified, and assembled with the linear parent PCR-amplified backbone using either PIPE or LIC cloning^87^, the latter of which incorporates an additional T4 polymerase digest of the insert and backbone mix prior to transformation. Plasmid libraries were electroporated into *E. coli* Dh5α to reach complete coverage (100,000-150,000 clones depending on library).

To confirm substitutions, 5 clones from each library were subjected to Sanger sequencing. Plasmid library colonies were scraped off of agar plates into LB medium containing 50 μg ml^−1^ kanamycin and normalized by colony number and OD to remove any biases that could skew the selection results using the following equation:

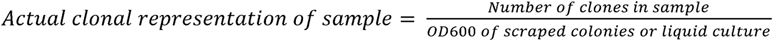

After combining, libraries were stored both as glycerol stocks and as plasmid preparations. RolR random variant libraries were constructed by error-prone PCR (epPCR) of the ligand-binding domain (residues 73-183). To generate epPCR libraries, a 50 μl PCR reaction was prepared by mixing the following: Taq buffer, 4mM MgCl_2_, 2 mM freshly prepared MnCl_2_, 1 μM each oligo, 1 μM CTP, 1 μM TTP, 0.2 μM ATP, 0.2 μM GTP, 20 ng template DNA, nuclease-free water, and Taq polymerase. For this library, we aimed for 1-2 amino acid substitutions per gene. To confirm the number of mutated residues, 3 clones from each library were subjected to Sanger sequencing. PCR products were gel-purified, PCR-amplified, and assembled with the linear parent PCR-amplified backbone using either PIPE or LIC cloning. Plasmid libraries were electroporated into *E. coli* Dh5α to reach a coverage of ~700,000 clones. Transformations were recovered in 1 ml LB medium for 1 h. After recovery, 50 uL were plated to identify the number of clones, and the remaining cells were pelleted and suspended in 5 ml LB medium containing 50 μg ml^−1^ kanamycin followed by incubating for 16 h at 37 °C. Cultures were normalized by colony number and OD using the same equation described above, combined, and stored both as glycerol stocks and as a plasmid preparation.

### Variant library selection

Dual selection was employed to simultaneously select for functional repressors that are capable of repressing DNA expression (negative selection), and for repressors that are induced by a ligand of interest (positive selection). To start dual selection, *E. coli* DH5α glycerol stocks containing the plasmid variant libraries were thawed, and 5 ml cultures were inoculated at a final OD_600_ of 0.05 into M9 medium containing 50 μg ml^−1^ kanamycin, and incubated for 16 h at 37 °C. For dual selection on solid media, which was implemented for the first-generation variant library, cells were plated on M9 agar plates containing 50 μg ml^−1^ kanamycin, 1 mM IPTG, and 0.6 mM NiCl_2_, and incubated at 37 °C for 48 h. For positive selection, colonies were scraped off, washed twice with M9 medium, plated on M9 agar plates containing 50 μg ml^−1^ kanamycin, 1 mM IPTG, 7.5 μg ml^−1^ tetracycline, and 0.5 mM catechol, and incubated for 16 h at 37 °C. This was followed by a final negative selection step for a total of 3 selection rounds (negative - positive - negative). The negative selection plates from the final round were subjected to an activity screening step described in the next section. Importantly, negative selection media components were mixed in the following order to avoid salt precipitation: NiCl_2_, CaCl_2_, MgSO_4_, followed by M9 salts. The rest of the media components (glucose, thiamine hydrochloride, casamino acids, kanamycin) were added in any order.

Alternatively, for liquid dual selection, which was implemented for the second and third variant library generations as they incorporated a larger number of small molecules, we followed the same procedure except for the addition of more selection rounds. Briefly, cells from overnight cultures were inoculated at a final OD_600_ of 0.05 into M9 medium containing 50 μg ml^−1^ kanamycin and 1 mM IPTG, and incubated for 16 h at 37 °C. For negative selection, cells were inoculated at a final OD_600_ of 0.005 into M9 medium containing 50 μg ml^−1^ kanamycin, 1 mM IPTG, and 0.4 mM NiCl_2_, and incubated at 37 °C for 16 hours. Cells were then inoculated at a final OD_600_ of 0.05 into M9 medium containing 50 μg ml^−1^ kanamycin, 1 mM IPTG, and 0.5 mM of the inducer of interest (catechol, protocatechuate, dopamine, L-DOPA, epinephrine, methyl catechol, caffeic acid, 2,3-dihydroxybenzoate, homovanillic acid, methoxytyramine), and incubated at 37 °C for 12 hours. Cells were then inoculated at a final OD_600_ of 0.005 into M9 medium containing 50 μg ml^−1^ kanamycin, 1 mM IPTG, and 7.5 μg ml^−1^ tetracycline, and 0.5 mM of the inducer of interest, and incubated at 37 °C for 12 hours. This process was repeated one more time ensuring that the last round is negative selection, for a total of 5 selection rounds (negative - positive - negative - positive - negative). Cells from the final liquid negative selection round were plated on LB plates containing 50 μg ml^−1^ kanamycin to obtain isolated colonies that were subjected to activity screening. Importantly, our selection scheme also included the addition of 50 mM of each molecule apart from the target for evolution in the negative selection step in an effort to minimize crosstalk, as higher concentrations were not possible due to the toxicity of these molecules.

### Biosensor activity screening

For variant characterization, ~480 colonies were picked from agar plates using a colony picker (Qpix 460) into five 96 well-plates with LB medium containing 50 μg ml^−1^ kanamycin, and were incubated for 16 h at 37 °C. The next day, 384-well plates were prepared for screening by dispensing 1 mM IPTG and 0.5 mM small molecules using an acoustic liquid handler (Labcyte Echo 550). This scheme will generate four data points per variant: an on state (no IPTG), an off state (IPTG), and 2 data points for testing inducibility by a small molecule of interest. Growing cells were diluted 1:100 into M9 medium containing 50 μg ml^−1^ kanamycin and transferred using a liquid handler (Biomek), and in some cases a 96-channel pipette (Gilson), into the 384 well-plates containing IPTG and small molecules, and plates were incubated for 20 h at 30 °C. The following day, eGFP fluorescence (excitation: 475-10 nm, emission: 510-10 nm) and OD_600_ of cultures were measured using a microplate reader (CLARIOstar, BMG Labtech, Germany). Data was processed using Excel, and positive hits that exhibited on, off, and small molecule-induced states were then subjected to two rounds of colony purification by streaking on LB plates containing 50 μg ml^−1^ kanamycin. Single colonies underwent a second activity screen, and positive hits were then subjected to Sanger sequencing.

### Biosensor characterization and data analyses

For biosensor response characterization, *E. coli* cells carrying genetic circuit plasmids were grown in LB medium containing 50 μg ml^−1^ kanamycin at 37 °C for 16 h. The following day, different volumes of IPTG and small molecules were dispensed into 96-well flat or deep-well using an acoustic liquid handler (Labcyte Echo 550) or an electronic multi-channel pipette (Thermo Scientific Finipipette), respectively. *E. coli* cells were diluted 1:100 in M9 medium containing 50 μg ml^−1^ kanamycin and 1mM IPTG, incubated at 37 °C for 4 h, and then diluted 1:100 in fresh M9 medium with the addition of small molecule ligands as needed, and incubated at 30 °C for 20 h. The following day, eGFP and OD_600_ of cultures were measured using a microplate reader (CLARIOstar, BMG Labtech, Germany) for first generation responses as well as dynamic range measurements for all variants, or using an Accuri C6 flow cytometer (BD Biosciences) for the second and third generation responses and crosstalk analyses. Cells were diluted 1:5 in deionized water to obtain an average rate of ~1000 events/second for a total of 10000 single-cell events. The mean fluorescence of the total ungated population was plotted for each molecule and/or concentration tested. Dose-response data points are averages of technical triplicates, with error bars representing standard deviations. When applicable, curves were fitted using GraphPad Prism by modelling them to a four-parameter non-linear regression equation, and EC_50_ values were determined as a result.

### Homology models and protein docking

When needed, homology models for variants were generated with Swiss-prot (https://swissmodel.expasy.org/) using RolR either in its apo (RCSB: 3AQS) or holo (RCSB: 3AQT) forms as a template. Models were then energy-minimized using the GROMOS96 force field tool of Swiss-PDB Viewer. Small molecule structures downloaded from PubChem or RCSB were docked into RolR variants using Autodock Vina^88^, and visualized using PyMol (DeLano Scientific). The lowest energy conformation with the ligand positioned in the designated binding pocket between helices 4,5,6 and 7 was used for further analyses.

### *Saccharomyces cerevisiae* strain construction

Biosensors were integrated in the genome of *S. cerevisiae* using CRISPR-Cas9-mediated genomic integration and *in vivo* DNA homologous recombination^89,90^. Briefly, yeast cells were transformed with a DNA pool comprising the following parts: PCR-amplified promoters, aTF, Envy GFP, and terminators with ~40 bp homology to the preceding and/or following part, mixed in equimolar ratios; two ~500 bp homology arms to the genomic locus targeted for integration (200 ng each); linearized pCas-G418 and pCas-Hyg vectors encoding Cas9^91^ (100 ng each) (Supplementary table 1); PCR-amplified gRNAs targeting the chromosomal locus Flagfeldt site 16^92^ and with homology to the Cas9 plasmid for assembly of the plasmid by *in vivo* homologous recombination (200 ng per transformation). The DNA pool was transformed into yeast using a 25 μL lithium acetate/PEG heat shock transformation^93^. Briefly, cells were heat-shocked at 42 °C for 30 min, recovered for 16 hours in YPD, and plated on YPD-Hyg/G418 dual selection plates. Successful genomic integrations were confirmed by colony PCR and Sanger sequencing. Finally, the Cas9 plasmid was cured by streaking colonies two times on YPD plates with no selection. Sequences of promoters, gene expression cassettes, terminators, and gRNAs are found in Supplementary data 2.

### *S. cerevisiae* biosensor characterization

Cells were grown overnight at 30 °C and 300 rpm in 96 deep-well plates in an orbital shaker. The following day, yeast cells were diluted 1:500 into 500 μL SC or YNB medium to an OD_600_ of ~0.01 in 96 deep-well plates with or without different concentrations of small molecules from the low μM range up to to 5-10 mM (depending on the toxicity of the molecule), and 100 mM in the case of protocatechuate. The plates were then incubated in an orbital shaker at 30 °C and 300 rpm for 20 h, after which fluorescence was measured using an Accuri C6 flow cytometer. Cells were diluted 1:5 in deionized water to obtain an average rate of ~1000 events/second for a total of 10000 single-cell events. The mean fluorescence of the total ungated population was plotted for each molecule and/or concentration tested. Unless otherwise stated, dose-response data points are averages of technical triplicates, shown either as 3 data points, or a single data point with an error bar representing standard deviation. When applicable, curves were fitted using GraphPad Prism by modelling them to a four-parameter nonlinear regression equation.

## Acknowledgements

This study was financially supported by NSERC Discovery Grants RGPIN-2016-05464 to D.H.K., and RGPIN-2017-06703 to V.J.J.M.; M.A.N. was supported by a Concordia International Tuition Award of Excellence, and an FRQNT B2X Doctoral Research Award. V.J.J.M. was supported by a Concordia University Research Chair. We thank Logan Timmins for contributing towards preliminary experiments. We thank Dr. Roberto Chica, Olivier Gagnon, and Dr. Anthony St-Jacques for their help in studying the structure of RolR, and with the choice of residues to target for mutagenesis. We thank the Concordia Genome Foundry for use of automation equipment, as well as James Bagley, Nicholas Gold, and Dr. Smita Amaranth for assistance with implementing and troubleshooting automation workflows.

## Author contributions

M.A.N., D.H.K., and V.J.J.M. designed the research. M.A.N. performed experiments and analyzed data. D.H.K. and V.J.J.M. supervised the research. M.A.N. wrote the manuscript with editing help from D.H.K., and V.J.J.M.

## Supplementary data

**Table 1:**
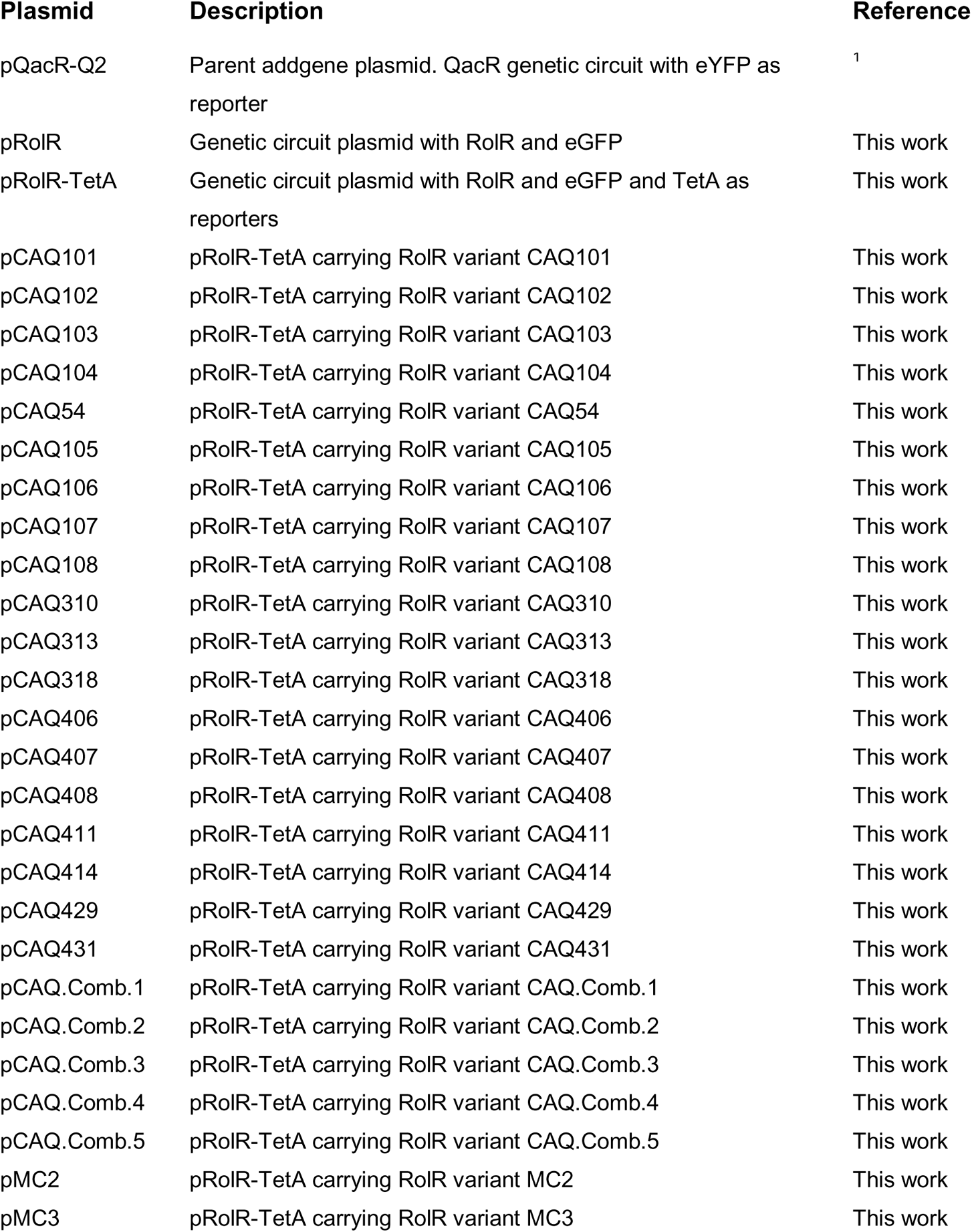
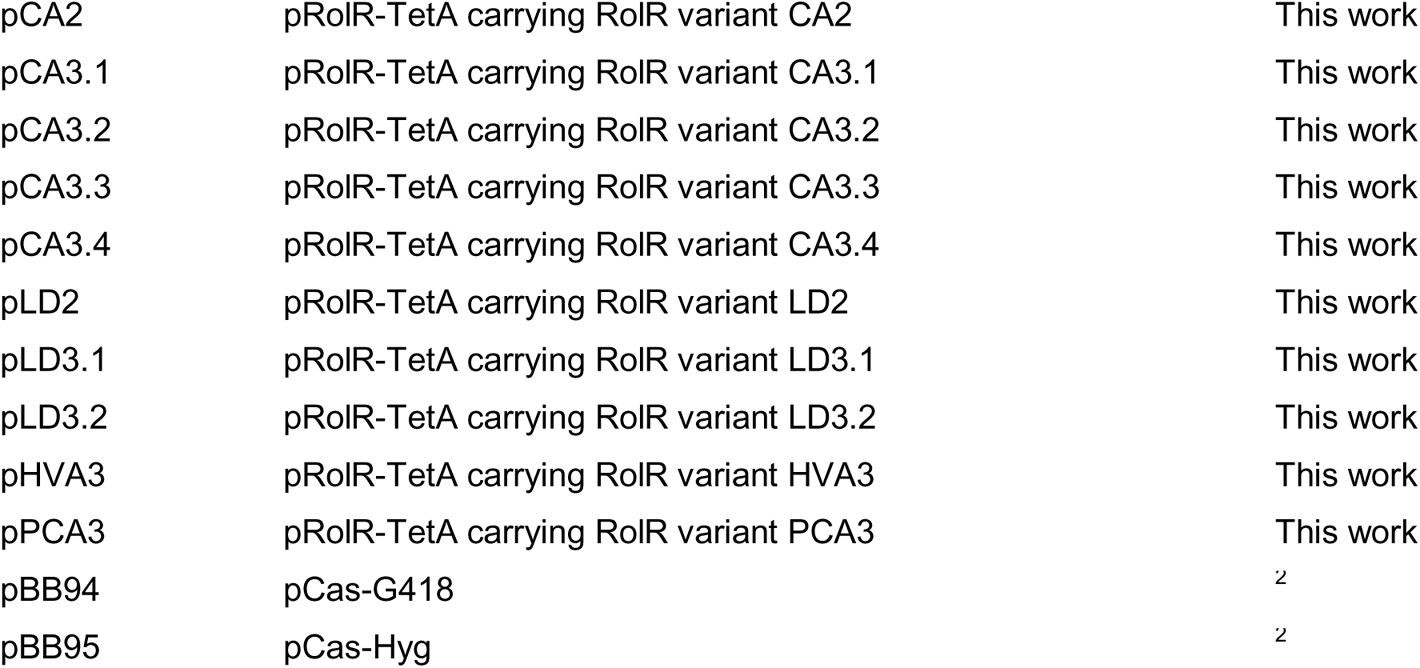
List of plasmids used in this study:

| Plasmid | Description | Reference |
| --- | --- | --- |
| pQacR-Q2 | Parent addgene plasmid. QacR genetic circuit with eYFP as reporter | <sup>1</sup> |
| pRoIR | Genetic circuit plasmid with RoIR and eGFP | This work |
| pRoIR-TetA | Genetic circuit plasmid with RoIR and eGFP and TetA as reporters | This work |
| pCAQ101 | pRoIR-TetA carrying RoIR variant CAQ101 | This work |
| pCAQ102 | pRoIR-TetA carrying RoIR variant CAQ102 | This work |
| pCAQ103 | pRoIR-TetA carrying RoIR variant CAQ103 | This work |
| pCAQ104 | pRoIR-TetA carrying RoIR variant CAQ104 | This work |
| pCAQ54 | pRoIR-TetA carrying RoIR variant CAQ54 | This work |
| pCAQ105 | pRoIR-TetA carrying RoIR variant CAQ105 | This work |
| pCAQ106 | pRoIR-TetA carrying RoIR variant CAQ106 | This work |
| pCAQ107 | pRoIR-TetA carrying RoIR variant CAQ107 | This work |
| pCAQ108 | pRoIR-TetA carrying RoIR variant CAQ108 | This work |
| pCAQ310 | pRoIR-TetA carrying RoIR variant CAQ310 | This work |
| pCAQ313 | pRoIR-TetA carrying RoIR variant CAQ313 | This work |
| pCAQ318 | pRoIR-TetA carrying RoIR variant CAQ318 | This work |
| pCAQ406 | pRoIR-TetA carrying RoIR variant CAQ406 | This work |
| pCAQ407 | pRoIR-TetA carrying RoIR variant CAQ407 | This work |
| pCAQ408 | pRoIR-TetA carrying RoIR variant CAQ408 | This work |
| pCAQ411 | pRoIR-TetA carrying RoIR variant CAQ411 | This work |
| pCAQ414 | pRoIR-TetA carrying RoIR variant CAQ414 | This work |
| pCAQ429 | pRoIR-TetA carrying RoIR variant CAQ429 | This work |
| pCAQ431 | pRoIR-TetA carrying RoIR variant CAQ431 | This work |
| pCAQ.Comb.1 | pRoIR-TetA carrying RoIR variant CAQ.Comb.1 | This work |
| pCAQ.Comb.2 | pRoIR-TetA carrying RoIR variant CAQ.Comb.2 | This work |
| pCAQ.Comb.3 | pRoIR-TetA carrying RoIR variant CAQ.Comb.3 | This work |
| pCAQ.Comb.4 | pRoIR-TetA carrying RoIR variant CAQ.Comb.4 | This work |
| pCAQ.Comb.5 | pRoIR-TetA carrying RoIR variant CAQ.Comb.5 | This work |
| pMC2 | pRoIR-TetA carrying RoIR variant MC2 | This work |
| pMC3 | pRoIR-TetA carrying RoIR variant MC3 | This work |
| pCA2 | pRoIR-TetA carrying RoIR variant CA2 | This work |
| pCA3.1 | pRoIR-TetA carrying RoIR variant CA3.1 | This work |
| pCA3.2 | pRoIR-TetA carrying RoIR variant CA3.2 | This work |
| pCA3.3 | pRoIR-TetA carrying RoIR variant CA3.3 | This work |
| pCA3.4 | pRoIR-TetA carrying RoIR variant CA3.4 | This work |
| pLD2 | pRoIR-TetA carrying RoIR variant LD2 | This work |
| pLD3.1 | pRoIR-TetA carrying RoIR variant LD3.1 | This work |
| pLD3.2 | pRoIR-TetA carrying RoIR variant LD3.2 | This work |
| pHVA3 | pRoIR-TetA carrying RoIR variant HVA3 | This work |
| pPCA3 | pRoIR-TetA carrying RoIR variant PCA3 | This work |
| pBB94 | pCas-G418 | <sup>2</sup> |
| pBB95 | pCas-Hyg | <sup>2</sup> |

**Supplementary table 2:**
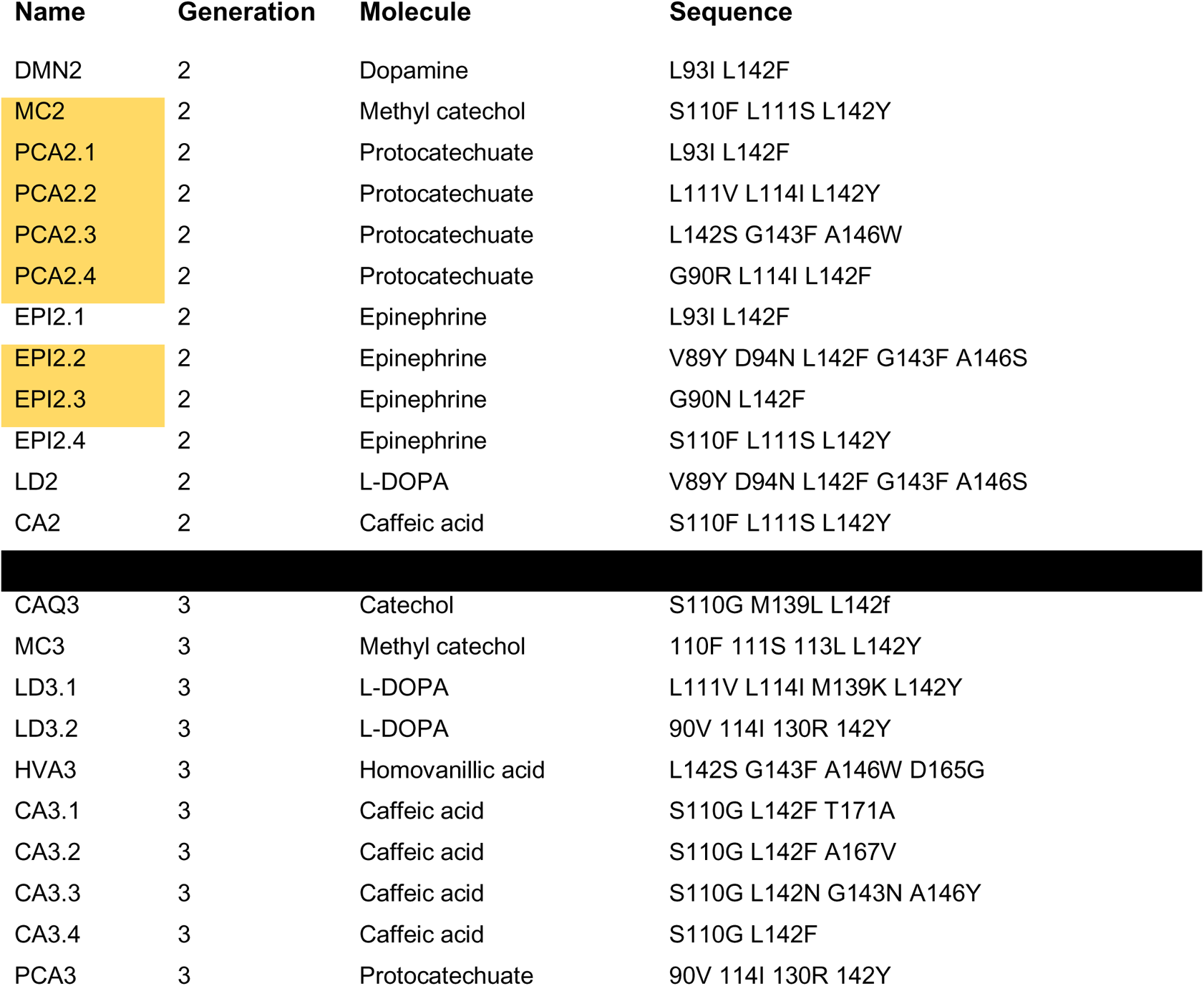
List of generation 2 and 3 RolR variants Sequences of variants: identified in the second and third generation with the molecules they were evolved towards. Second generation variants highlighted in yellow were used as a template for random mutagenesis, yielding generation 3 variants.

**Supplementary Figure 1.**
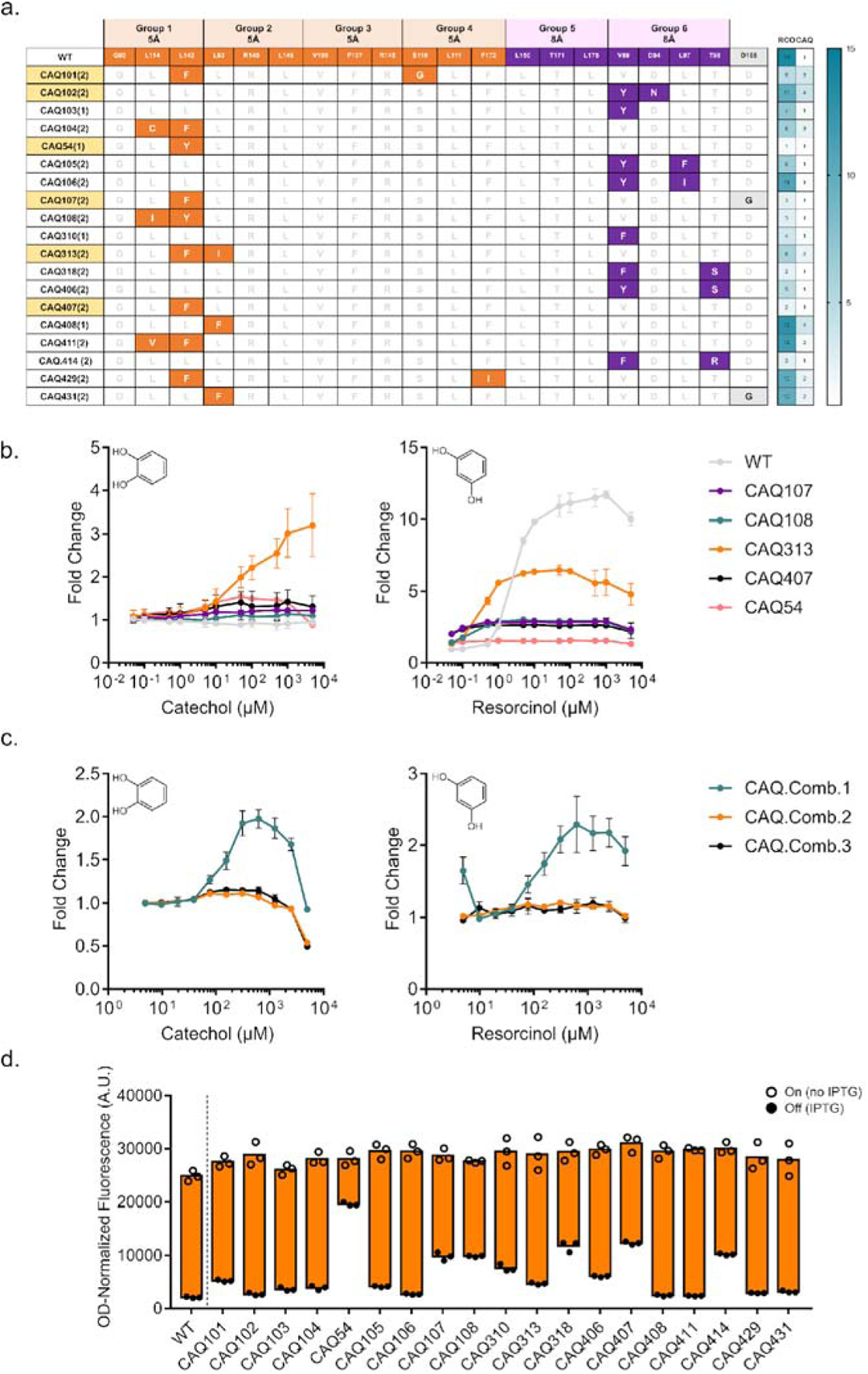
Continued first generation variant hits towards catechol. **a** Table depicting identified variants responsive to catechol, their substitutions coloured according to group, and their response towards resorcinol and catechol. **b** Dose response curves of the remaining variants compared to WT RolR with catechol (left panel) and resorcinol (right panel). **c** Dose response curves of the combinatorial variants with resorcinol (left panel) and catechol (right panel). **d** Dynamic range of WT RolR and all identified variants.

**Supplementary Figure 2.**
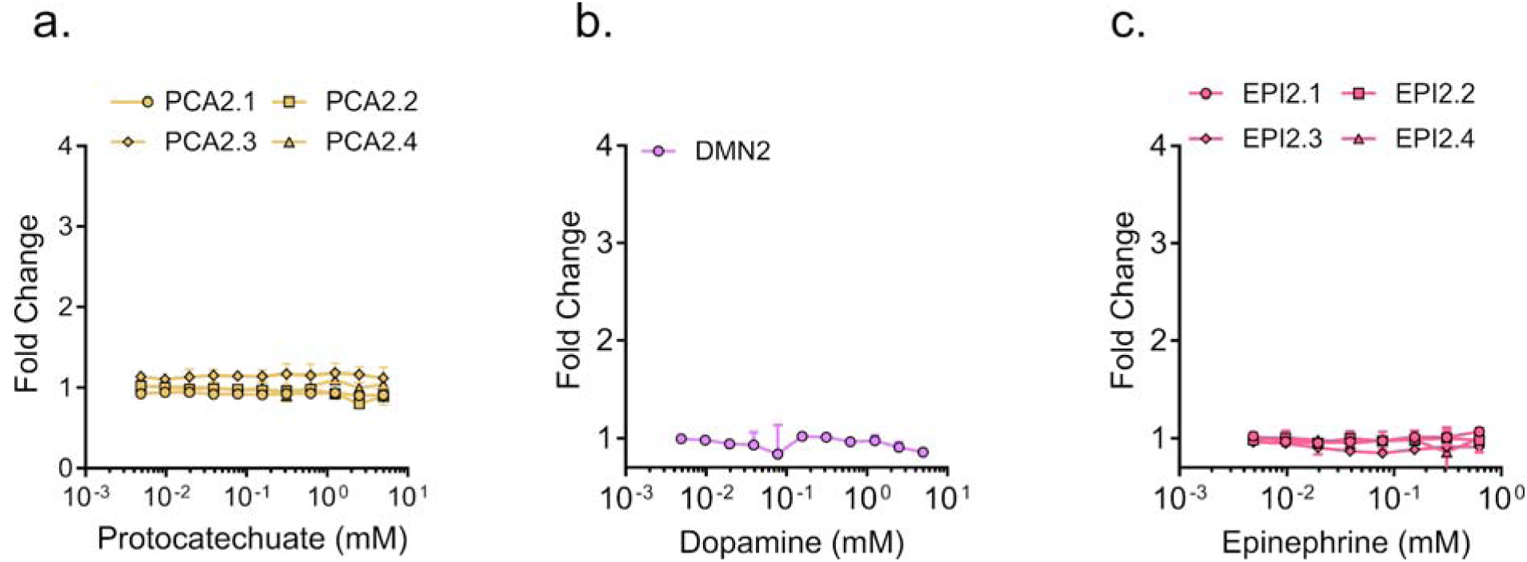
Dose response curves for identified second generation variant(s). Dose response curves for variants identified in the second generation of mutagenesis towards protocatechuate **(a)**, dopamine **(b)**, and epinephrine **(c)**.

**Supplementary Figure 3.**
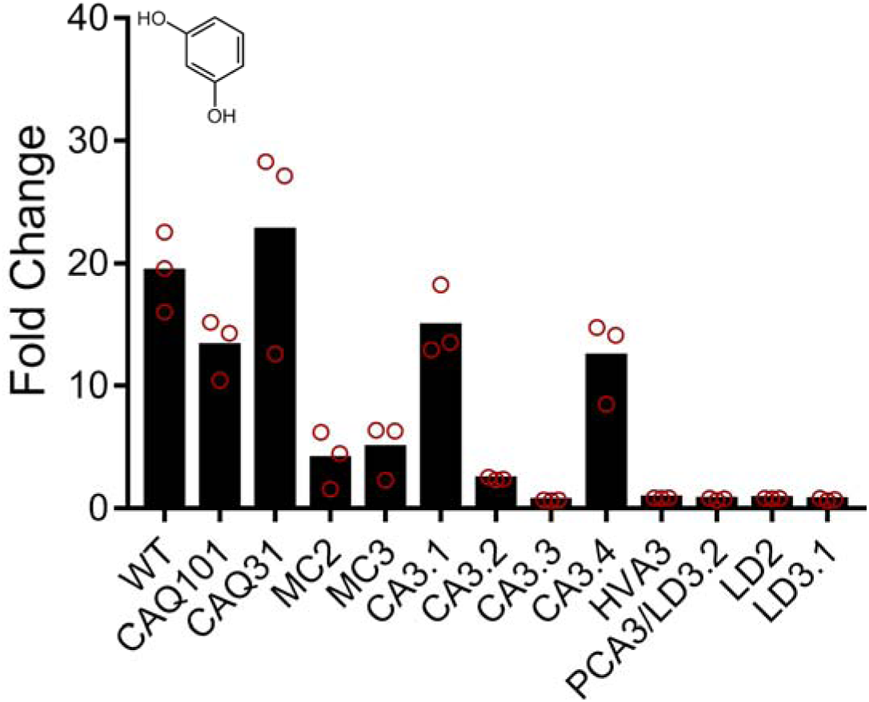
RolR variant response to resorcinol. Fluorescence fold-change response of RolR variants to resorcinol.

**Supplementary Figure 4.**
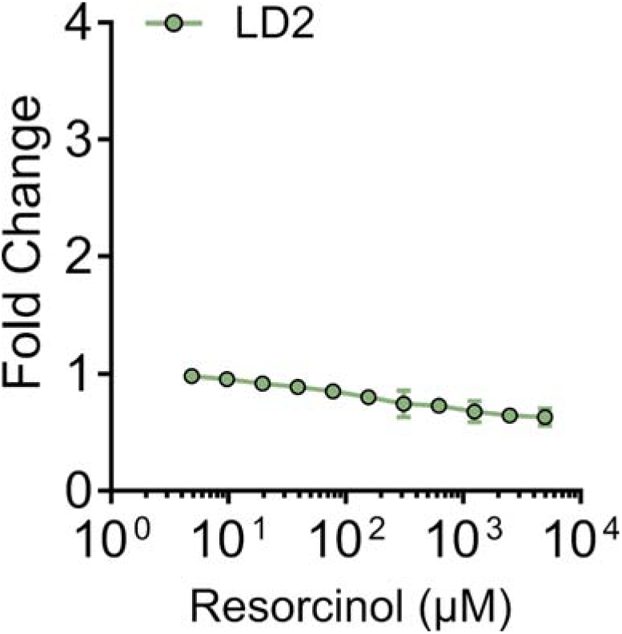
Dose response curves for identified second generation variant(s). Dose response curve for variant LD2 with resorcinol.

**Supplementary Figure 5.**
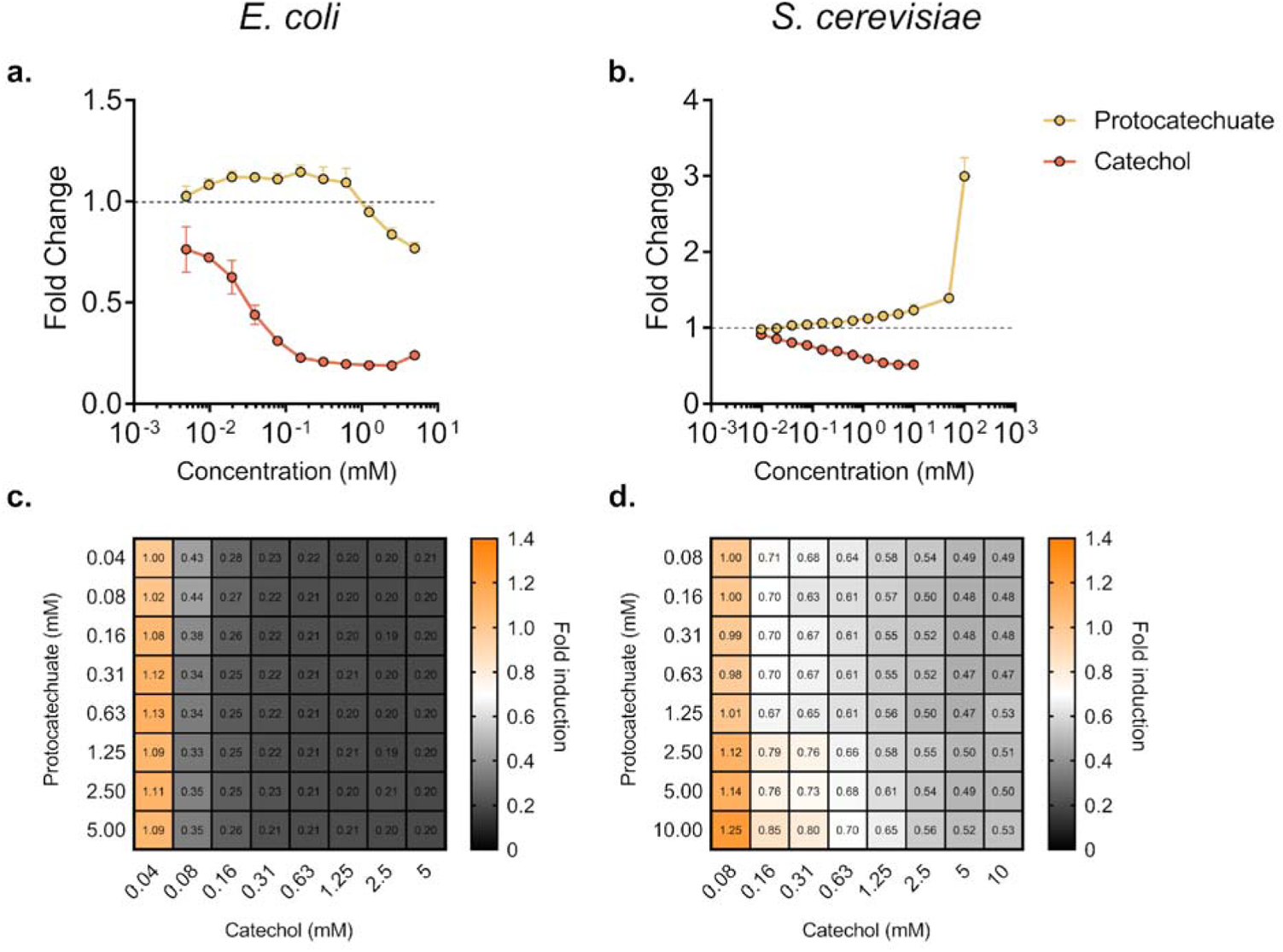
Engineering a tri-stable aTF with function towards catechol and protocatechuate. Dose response curve of sensor PCA3 with catechol superimposed with that of protocatechuate in *E. coli* **(a)** and *S. cerevisiae* **(b)**. GFP fluorescence after titrating different concentrations of catechol and protocatechuate in *E. coli* **(c)** and *S. cerevisiae* **(d)**.

