## Supplementary Table 1, Supplementary Table 2 for "Divergent Directed Evolution of a TetR-type Repressor Towards Aromatic Molecules"

**Supplementary table 1: List of plasmids used in this study:**

| **Plasmid** | **Description** | **Reference** |
| --- | --- | --- |
| pQacR-Q2 | Parent addgene plasmid. QacR genetic circuit with eYFP as reporter | ^1^ |
| pRolR | Genetic circuit plasmid with RolR and eGFP | This work |
| pRolR-TetA | Genetic circuit plasmid with RolR and eGFP and TetA as reporters | This work |
| pCAQ101 | pRolR-TetA carrying RolR mutant CAQ101 | This work |
| pCAQ102 | pRolR-TetA carrying RolR mutant CAQ102 | This work |
| pCAQ103 | pRolR-TetA carrying RolR mutant CAQ103 | This work |
| pCAQ104 | pRolR-TetA carrying RolR mutant CAQ104 | This work |
| pCAQ54 | pRolR-TetA carrying RolR mutant CAQ54 | This work |
| pCAQ105 | pRolR-TetA carrying RolR mutant CAQ105 | This work |
| pCAQ106 | pRolR-TetA carrying RolR mutant CAQ106 | This work |
| pCAQ107 | pRolR-TetA carrying RolR mutant CAQ107 | This work |
| pCAQ108 | pRolR-TetA carrying RolR mutant CAQ108 | This work |
| pCAQ310 | pRolR-TetA carrying RolR mutant CAQ310 | This work |
| pCAQ313 | pRolR-TetA carrying RolR mutant CAQ313 | This work |
| pCAQ318 | pRolR-TetA carrying RolR mutant CAQ318 | This work |
| pCAQ406 | pRolR-TetA carrying RolR mutant CAQ406 | This work |
| pCAQ407 | pRolR-TetA carrying RolR mutant CAQ407 | This work |
| pCAQ408 | pRolR-TetA carrying RolR mutant CAQ408 | This work |
| pCAQ411 | pRolR-TetA carrying RolR mutant CAQ411 | This work |
| pCAQ414 | pRolR-TetA carrying RolR mutant CAQ414 | This work |
| pCAQ429 | pRolR-TetA carrying RolR mutant CAQ429 | This work |
| pCAQ431 | pRolR-TetA carrying RolR mutant CAQ431 | This work |
| pCAQ.Comb.1 | pRolR-TetA carrying RolR mutant CAQ.Comb.1 | This work |
| pCAQ.Comb.2 | pRolR-TetA carrying RolR mutant CAQ.Comb.2 | This work |
| pCAQ.Comb.3 | pRolR-TetA carrying RolR mutant CAQ.Comb.3 | This work |
| pCAQ.Comb.4 | pRolR-TetA carrying RolR mutant CAQ.Comb.4 | This work |
| pCAQ.Comb.5 | pRolR-TetA carrying RolR mutant CAQ.Comb.5 | This work |
| pMC2 | pRolR-TetA carrying RolR mutant MC2 | This work |
| pMC3 | pRolR-TetA carrying RolR mutant MC3 | This work |
| pCA2 | pRolR-TetA carrying RolR mutant CA2 | This work |
| pCA3.1 | pRolR-TetA carrying RolR mutant CA3.1 | This work |
| pCA3.2 | pRolR-TetA carrying RolR mutant CA3.2 | This work |
| pCA3.3 | pRolR-TetA carrying RolR mutant CA3.3 | This work |
| pCA3.4 | pRolR-TetA carrying RolR mutant CA3.4 | This work |
| pLD2 | pRolR-TetA carrying RolR mutant LD2 | This work |
| pLD3.1 | pRolR-TetA carrying RolR mutant LD3.1 | This work |
| pLD3.2 | pRolR-TetA carrying RolR mutant LD3.2 | This work |
| pHMV3 | pRolR-TetA carrying RolR mutant HMV3 | This work |
| pPCA3 | pRolR-TetA carrying RolR mutant PCA3 | This work |
| pBB94 | pCAS-G418 | ^2^ |
| pBB95 | pCAS-Hyg | ^2^ |

**Supplementary table 2: List of generation 2 and 3 RolR mutants:**

| **Name** | **Generation** | **Molecule** | **Sequence** |
| --- | --- | --- | --- |
| DMN2 | 2 | Dopamine | L93I L142F |
| MC2 | 2 | Methyl catechol | S110F L111S L142Y |
| PCA2.1 | 2 | Protocatechuate | L93I L142F |
| PCA2.2 | 2 | Protocatechuate | L111V L114I L142Y |
| PCA2.3 | 2 | Protocatechuate | L142S G143F A146W |
| PCA2.4 | 2 | Protocatechuate | G90R L114I L142F |
| EPI2.1 | 2 | Epinephrine | L93I L142F |
| EPI2.2 | 2 | Epinephrine | V89Y D94N L142F G143F A146S |
| EPI2.3 | 2 | Epinephrine | G90N L142F |
| EPI2.4 | 2 | Epinephrine | S110F L111S L142Y |
| LD2 | 2 | L-DOPA | V89Y D94N L142F G143F A146S |
| CA2 | 2 | Caffeic acid | S110F L111S L142Y |
| CAQ3 | 3 | Catechol | S110G M139L L142f |
| MC3 | 3 | Methyl catechol | 110F 111S 113L L142Y |
| LD3.1 | 3 | L-DOPA | L111V L114I M139K L142Y |
| LD3.2 | 3 | L-DOPA | 90V 114I 130R 142Y |
| HMV3 | 3 | Homovanillic acid | L142S G143F A146W D165G |
| CA3.1 | 3 | Caffeic acid | S110G L142F T171A |
| CA3.2 | 3 | Caffeic acid | S110G L142F A167V |
| CA3.3 | 3 | Caffeic acid | S110G L142N G143N A146Y |
| CA3.4 | 3 | Caffeic acid | S110G L142F |
| PCA3 | 3 | Protocatechuate | 90V 114I 130R 142Y |

**Supplementary table 2** Sequences of mutants identified in the second and third generation with the molecules they were evolved towards. Second generation mutants highlighted in yellow were used as a template for random mutagenesis, yielding generation 3 mutants.

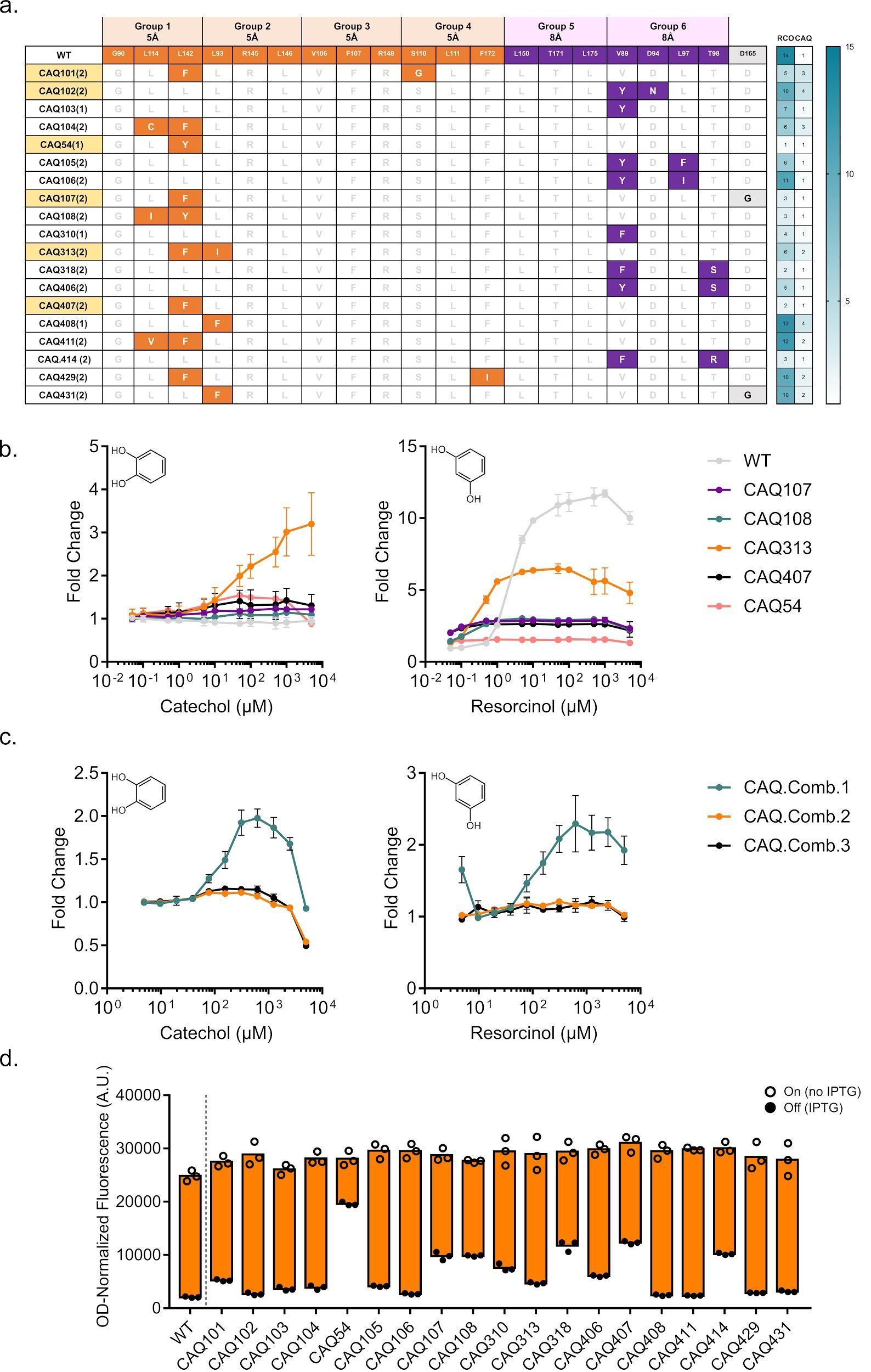

**Supplementary Figure 1.** **Continued** **first generation variant hits towards catechol*.* a** Table depicting identified variants responsive to catechol, their substitutions coloured according to group, and their response towards resorcinol and catechol. **b** Dose response curves of the remaining variants compared to WT RolR with catechol (left panel) and resorcinol (right panel). **c** Dose response curves of the combinatorial variants with resorcinol (left panel) and catechol (right panel). **d** Dynamic range of WT RolR and all identified variants.

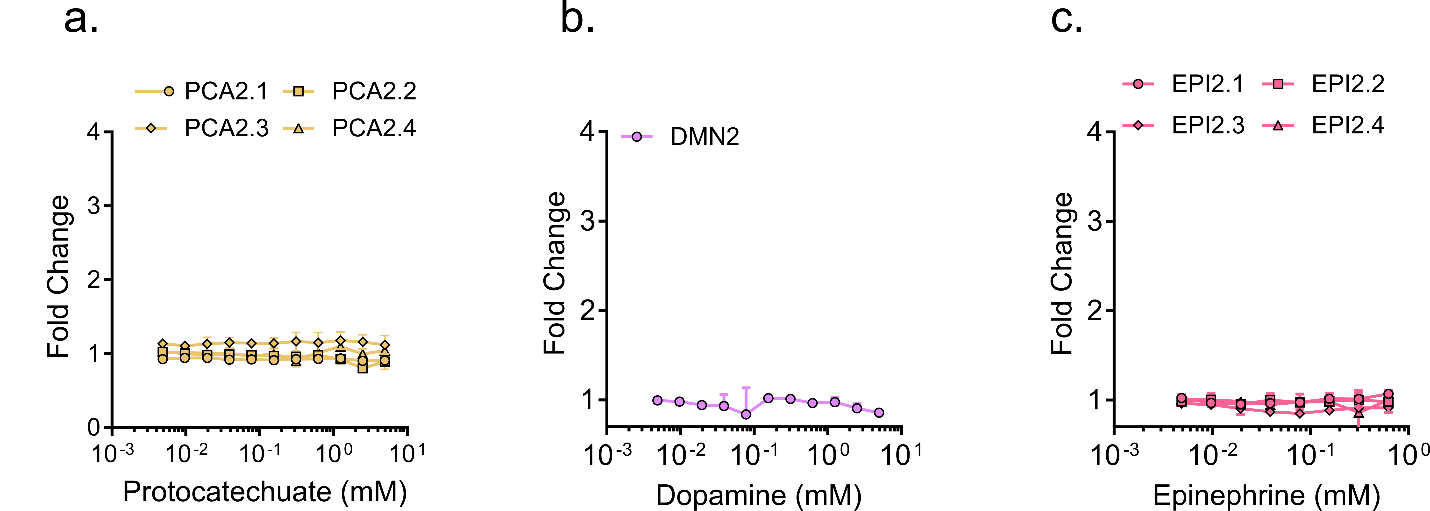

**Supplementary Figure 2. Dose response curves for identified second generation variant(s).** Dose response curves for variants identified in the second generation of mutagenesis towards protocatechuate **(a)**, dopamine **(b)**, and epinephrine **(c)**.

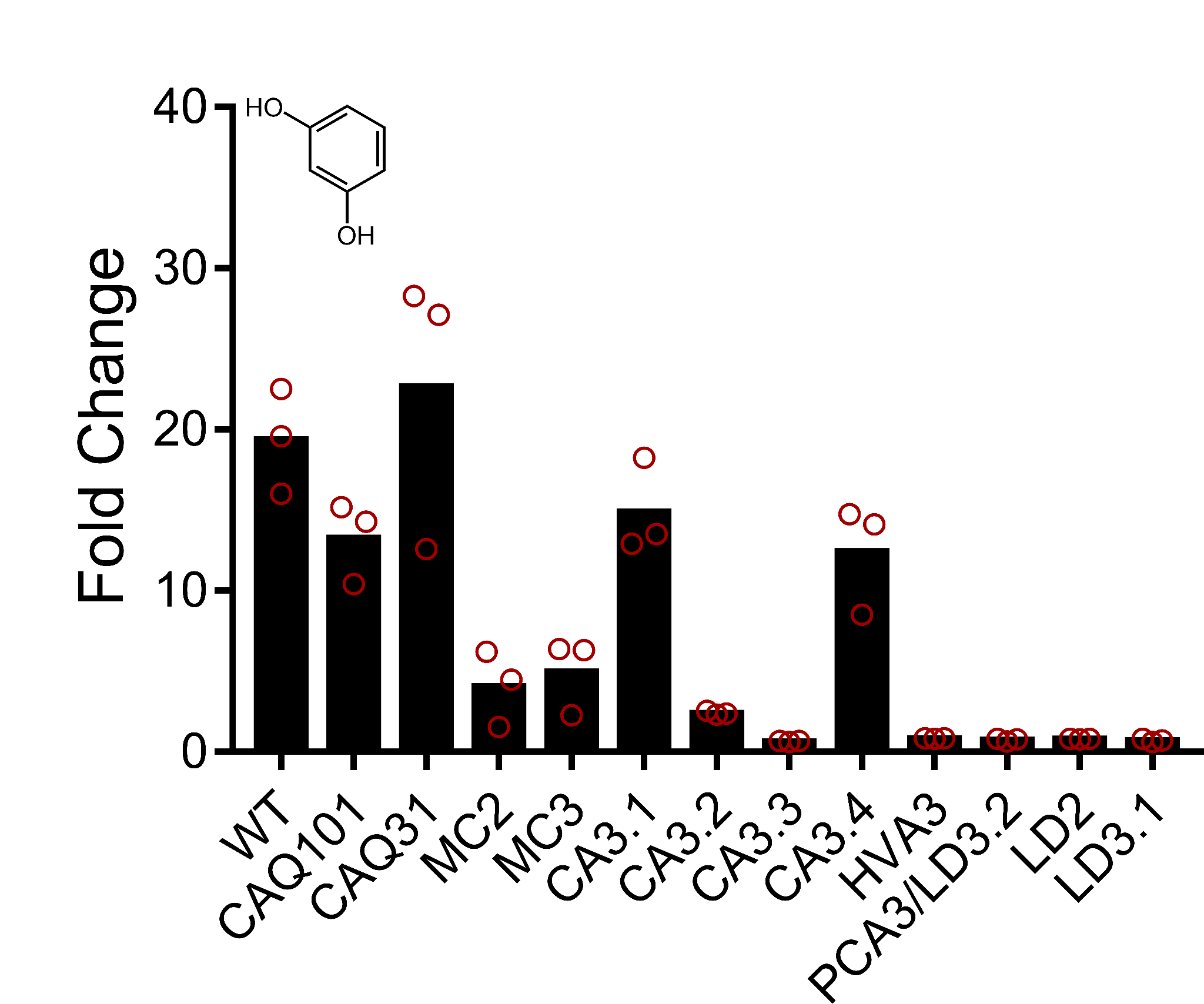

**Supplementary Figure 3. RolR variant response to resorcinol.** Fluorescence fold-change response of RolR variants to resorcinol.

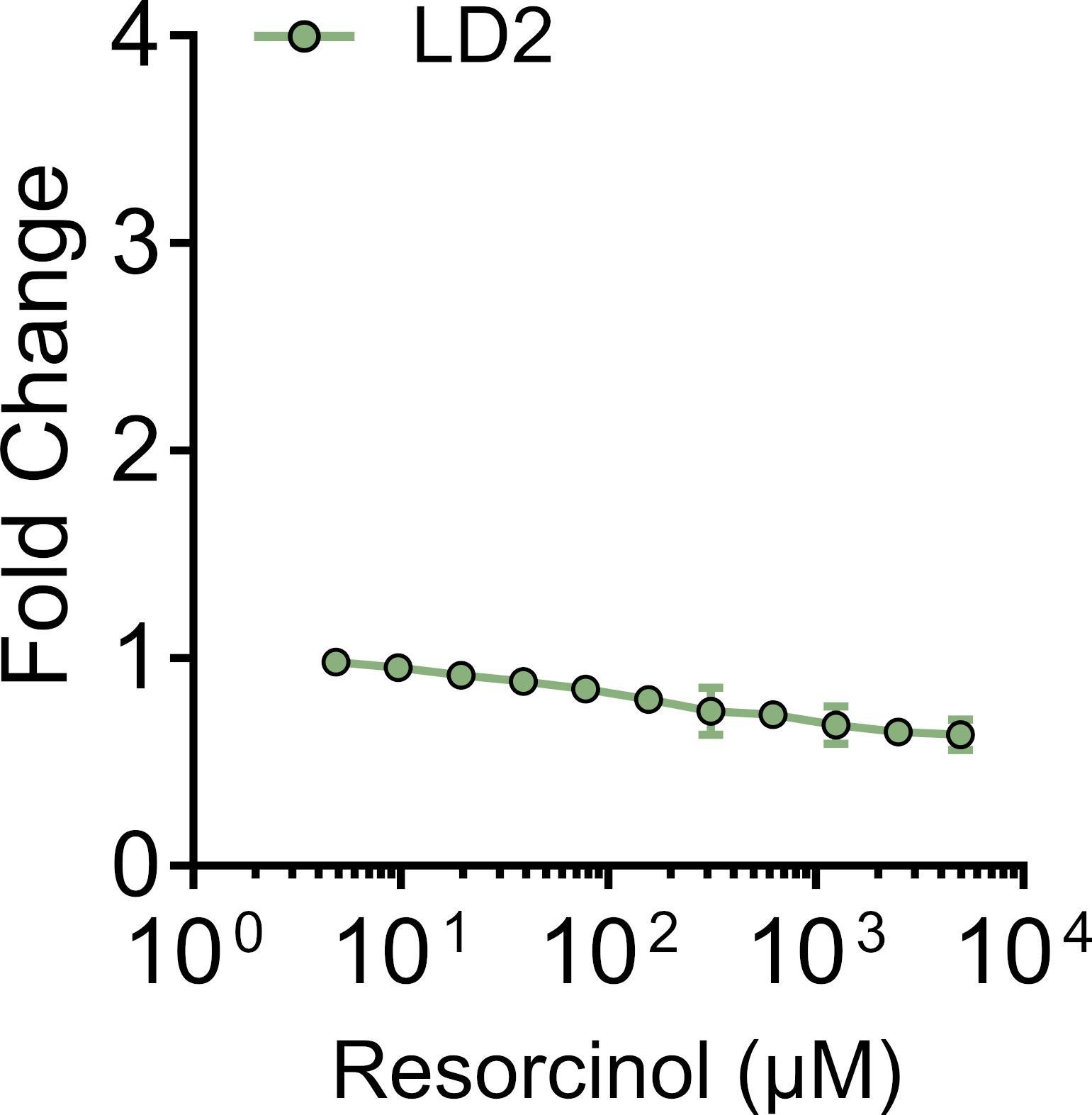

**Supplementary Figure 4. Dose response curves for identified second generation variant(s).**

Dose response curve for variant LD2 with resorcinol.

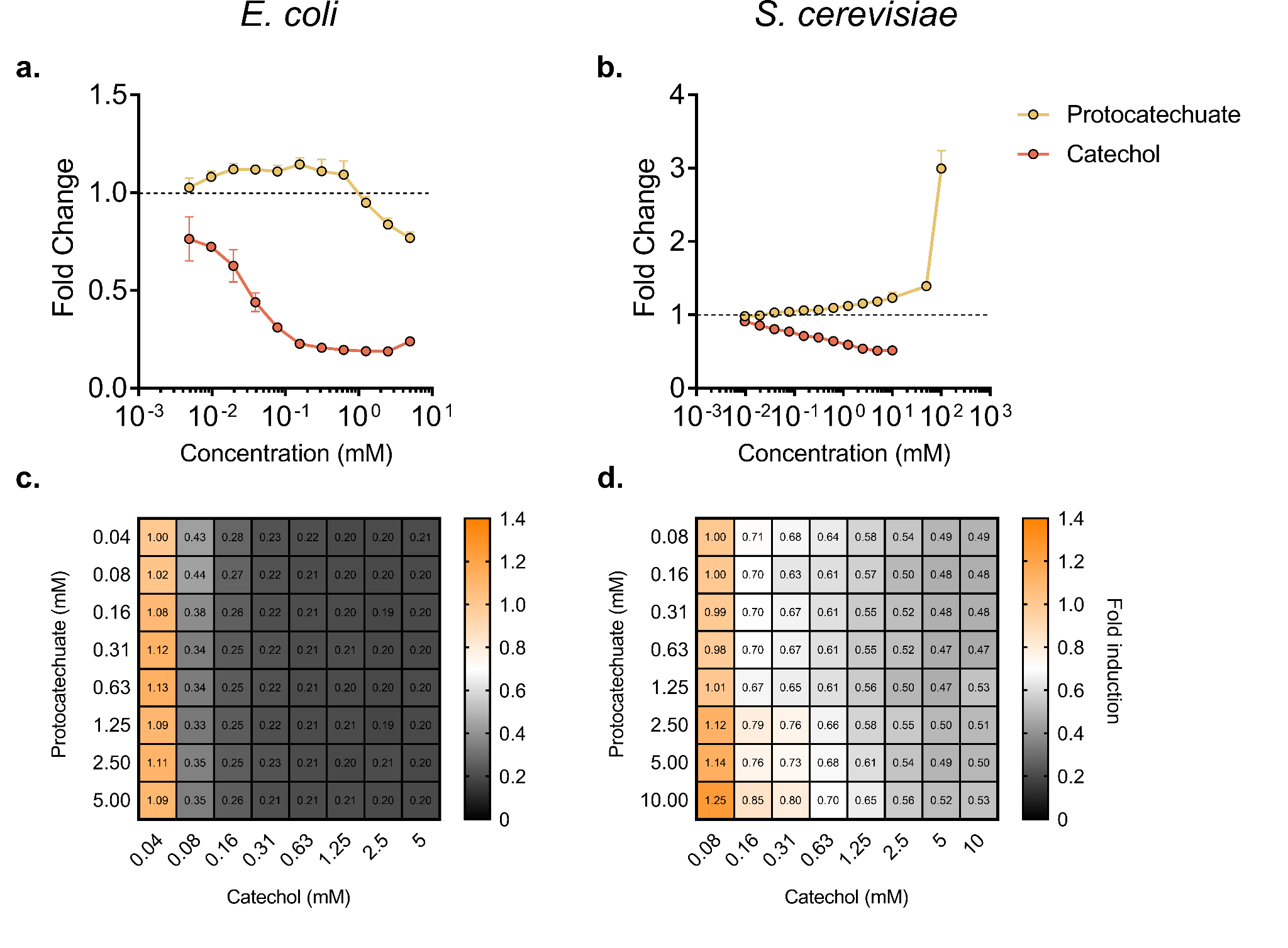

**Supplementary Figure 5. Engineering a tri-stable aTF with function towards catechol and protocatechuate.** Dose response curve of sensor PCA3 with catechol superimposed with that of protocatechuate in *E. coli* **(a)** and *S. cerevisiae* **(b)**. GFP fluorescence after titrating different concentrations of catechol and protocatechuate in *E. coli* **(c)** and *S. cerevisiae* **(d)**.
