## Supplementary figures and images for "Divergent Directed Evolution of a TetR-type Repressor Towards Aromatic Molecules"

### Supplementary Fig. 1

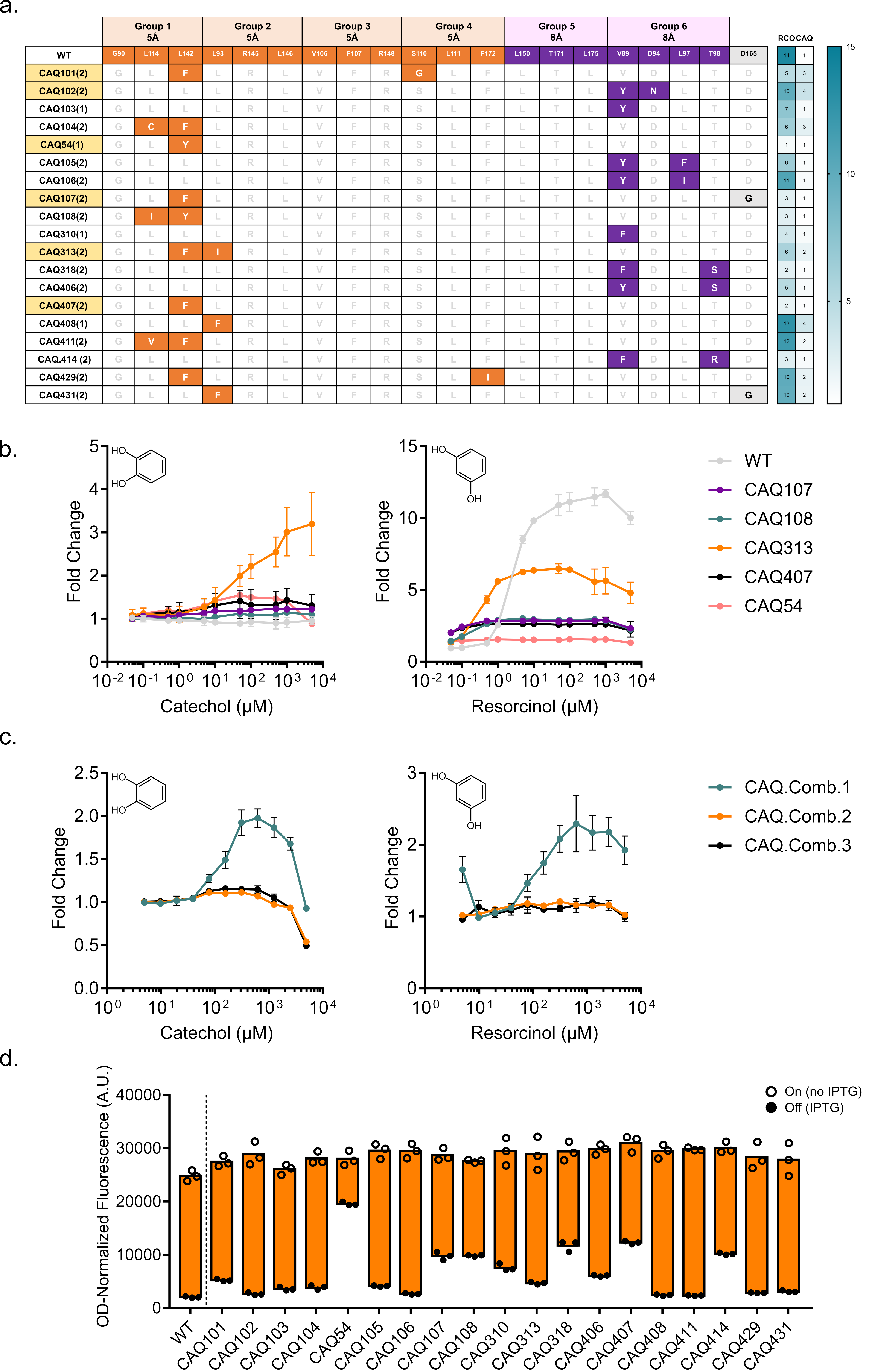

### Supplementary Fig. 2

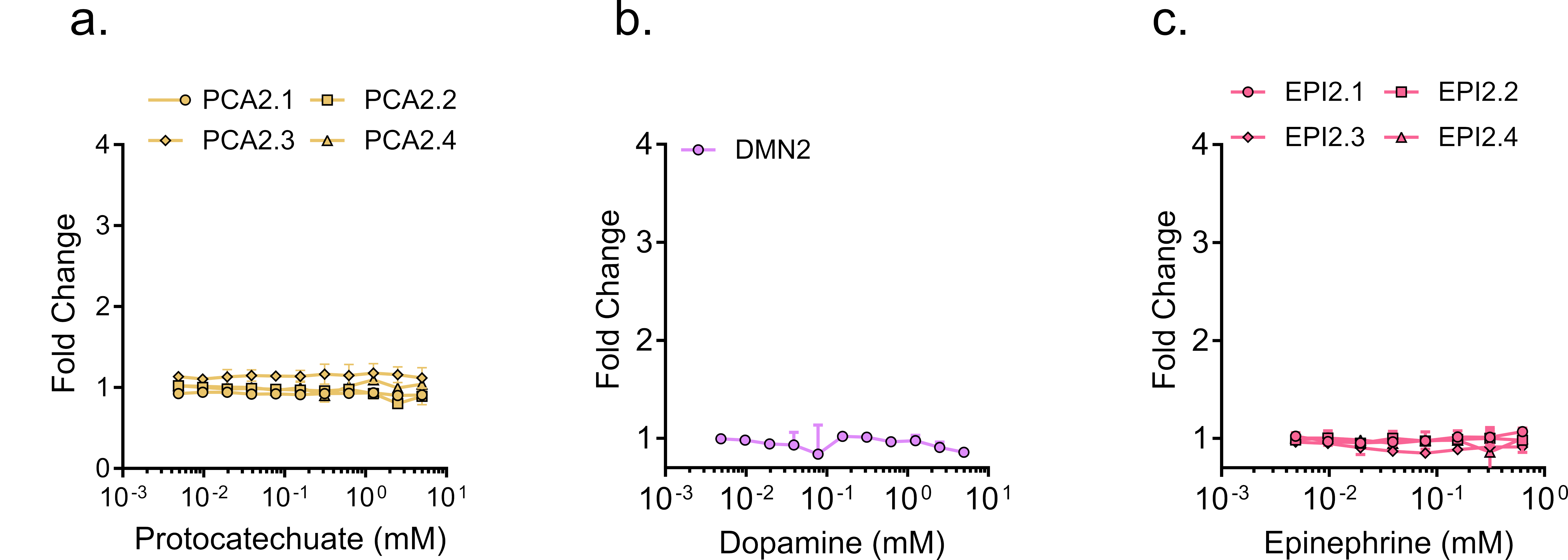

### Supplementary Fig. 3

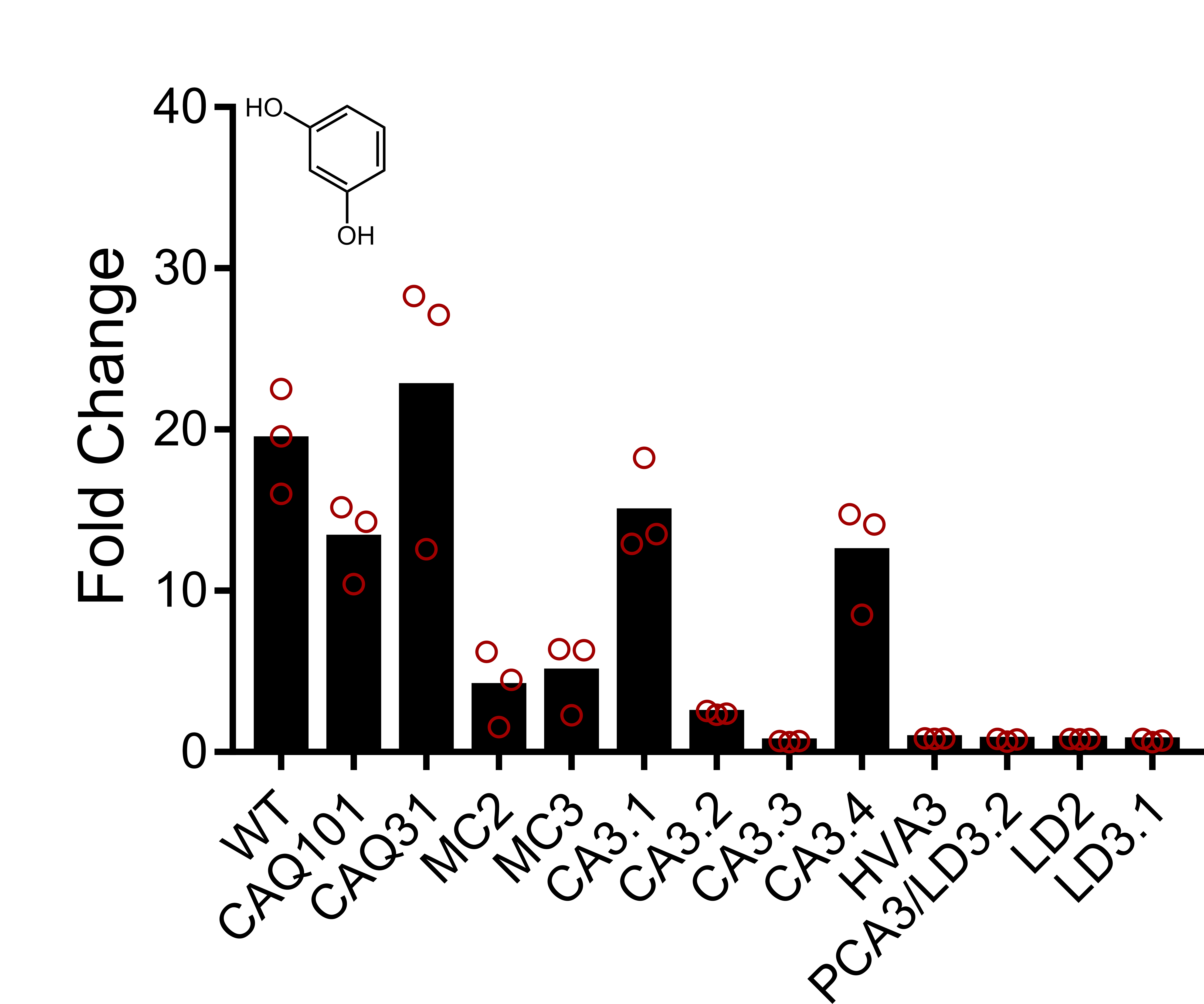

### Supplementary Fig. 4

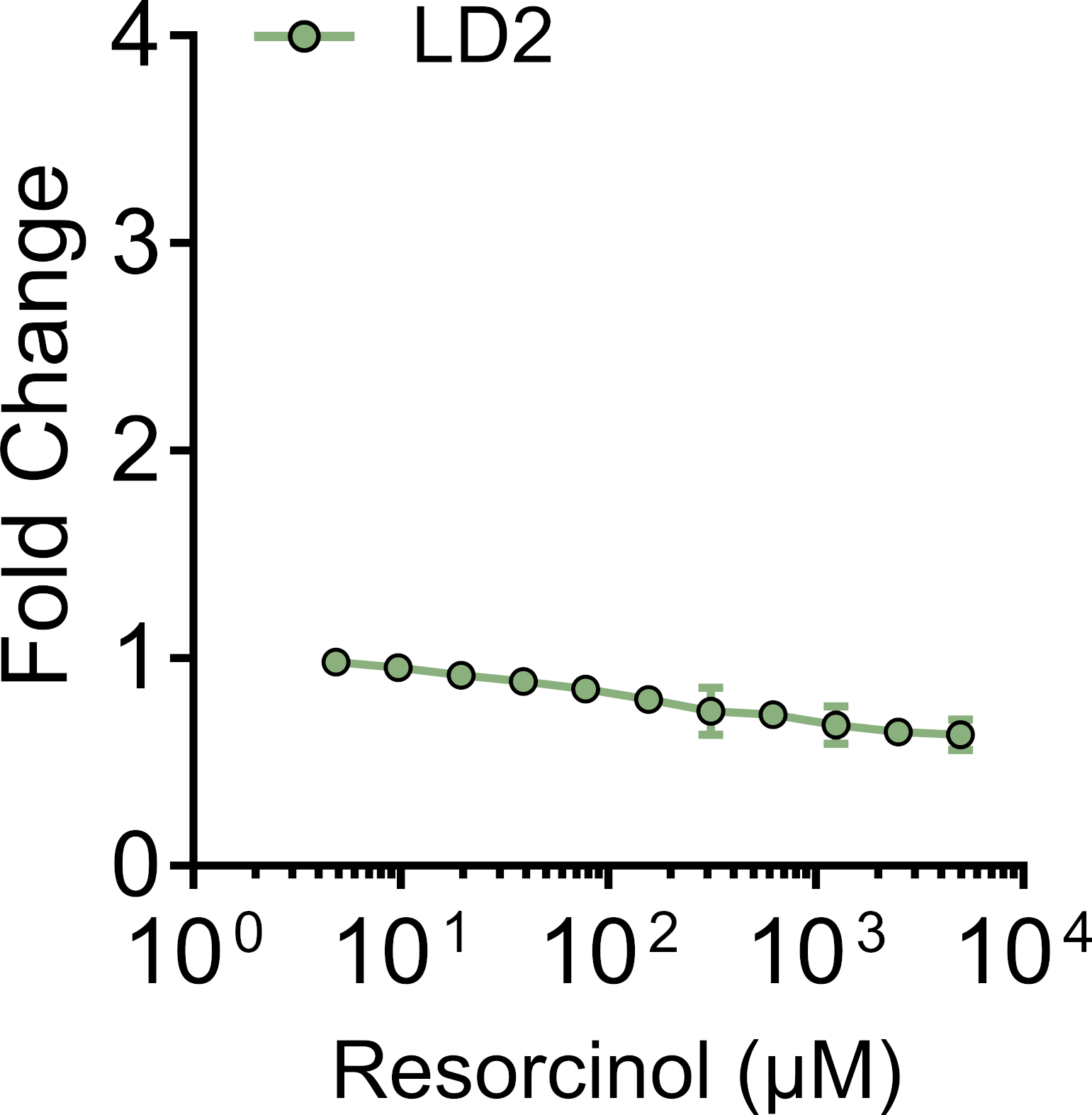

### Supplementary Fig. 5

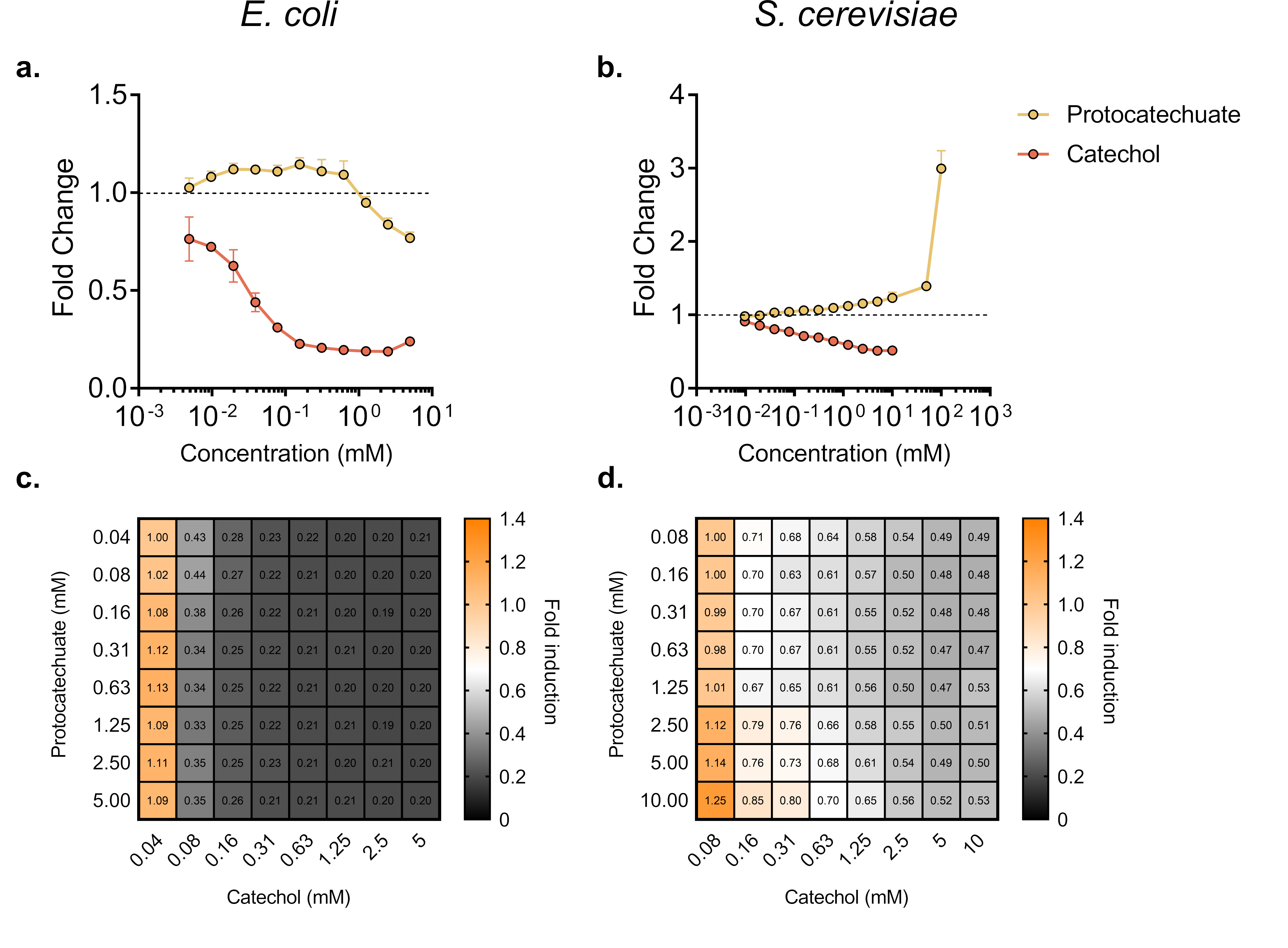
